# Precise spatiotemporal control of voltage-gated sodium channels by photocaged saxitoxin

**DOI:** 10.1101/2020.11.18.389130

**Authors:** Anna V. Elleman, Gabrielle Devienne, Christopher D. Makinson, Allison L. Haynes, John R. Huguenard, J. Du Bois

**Author notes:** **Corresponding Authors**, Correspondence to J. Du Bois or John R. Huguenard. Institute for Genomic Medicine, Columbia University Irving Medical Center, New York NY USA; Department of Neurology, Columbia University Irving Medical Center, New York, NY USA.

## Abstract

Here we report the pharmacologic blockade of voltage-gated sodium ion channels (NaV) by a synthetic saxitoxin derivative affixed to a photocleavable protecting group. We demonstrate that a functionalized saxitoxin (STX-eac) enables exquisite spatiotemporal control of NaV blockade to interrupt action potentials (APs) in dissociated neurons and nerve fiber bundles. The photo-uncaged inhibitor (STX-ea) is a nanomolar potent, reversible binder of NaVs. We use STX-eac to reveal differential susceptibility of myelinated and unmyelinated axons in the corpus callosum to NaV-dependent alterations in AP propagation, with unmyelinated axons preferentially showing reduced AP fidelity under conditions of partial NaV blockade. These results validate STX-eac as a high precision tool for robust photocontrol of neuronal excitability and AP generation.

## Introduction

Small molecules bearing photocleavable protecting groups^1^ have been used extensively to manipulate biochemical processes, including protein binding interactions ^2, 3, 4^, receptor activation^5^, and signaling pathway dynamics^6,7^. Photocaged molecules provide unique advantages over traditional pharmacologic agents and pro-drugs, namely precise temporal and spatial release, tunability, and reagent-less deprotection. Neuroactive compounds including GABA^8^, glutamic acid^9,10^, dopamine^11^, serotonin^12^, Leu-enkephalin^13^, cAMP^14^, lipids^15^, and calcium^16^ have been photocaged in order to study signaling mechanisms in cells, tissues, and whole organisms. Despite this impressive collection of photo-protected reagents, to our knowledge, no such compound has been designed to directly target action potentials (APs), the fundamental mode of communication in electrically excitable cells. Herein, we demonstrate that photochemical release of a caged form of saxitoxin (STX-eac) impedes AP initiation and propagation by blocking voltage-gated sodium ion channel (NaV) function. Experiments with dissociated neuronal cells and in tissue slice validate the utility of this unique tool compound for studies of electrical signaling.

Modern methods in neuroscience enable precise optical and chemical control of neuronal activity. Viral transduction of optogenetic and chemogenetic ion channels and receptors makes possible specific activation or inactivation of defined cell populations^17,18,19,20^. These technologies are neuromodulating (i.e., increase or decrease signaling likelihood) and generally do not completely silence neuronal outputs^21,22^. NaVs are responsible for mediating AP initiation and propagation and are therefore an ideal target for controlling electrical transmission. Furthermore, the location, amplitude, frequency, and timing of sodium ion influx are important factors that determine electrical excitability, synaptic release, and ultimately information flow through neural systems^23,24^.

In addition to affecting AP properties, a growing body of evidence implicates NaV activity in modulating dendritic currents, synaptic release, homeostasis, circuit function, and ultimately, behavior^25^. Studies of NaV physiology have primarily relied on pharmacological blockade of axon excitability using partially inactivating concentrations^26,27^ or local pressure puffing of small molecule inhibitors^28,29^; however, these methods do not provide the fine spatial or temporal resolution required to specifically inactivate selected cells or NaVs in cellular compartments. The design and development of photocaged NaV inhibitors for precise control of neuronal membrane excitability addresses these limitations.

### Synthesis of photocaged STX derivatives

In order to achieve spatiotemporal control of NaVs, a photocaging group was appended to a synthetic derivative of saxitoxin (STX), a naturally occurring bis-guanidinium toxin that specifically inhibits the action of six of nine NaV subtypes (e.g., IC_50_ = 1.2 nM vs. rat NaV 1.2)^30^. STX targets the outer mouth of the NaV pore, thereby occluding ion passage into cells. The location of the STX binding site enables extracellular application, and block by this small molecule toxin is entirely reversible, thus advantaging it over other NaV antagonists. Modification of the N21 carbamate of STX affords saxitoxin ethylamine (STX-ea **1**)^31^, a toxin derivative of similar potency to that of the natural product (IC_50_ = 14.4 nM vs. rat NaV1.2). Different prosthetic groups, including photolabile carbamate derivatives, can be attached to the primary amine in STX-ea through selective acylation reactions. This tactic allows for introduction of the photocage in the final step of the synthetic sequence and is considerably more efficient than protection of either or both of the guanidinium groups. We exploited this chemistry to access four coumarin-derived photocaged STXs, including STX-eac **5**(**Figure 1A**).

**Fig. 1:**
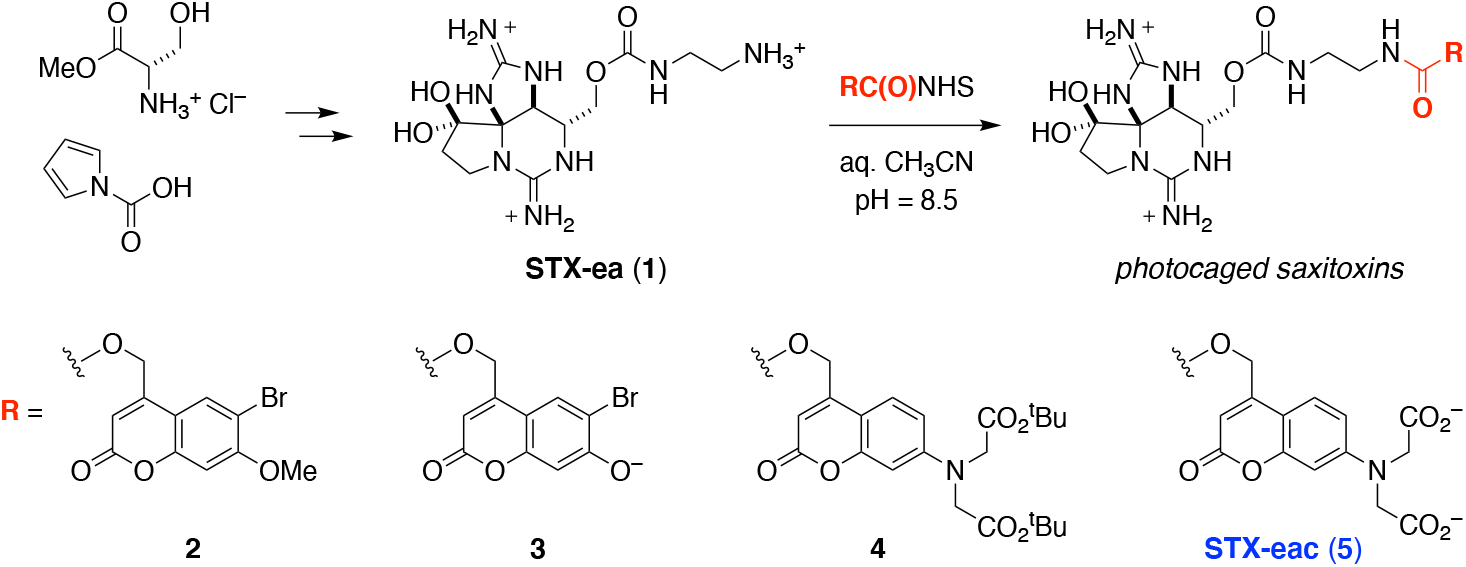
Caging STX-ea **1**with photo-protecting groups. (**A**) Synthesis and selective carbamoylation of STX-ea **1**. (**B**) Photocaged derivatives of STX-ea.

Coumarin photo-protecting groups are well-established and display excellent one- and two-photon uncaging efficiencies^32,33^. Additionally, the byproducts of uncaging are generally inert to biological tissue^34^ (**Figure 1B**). Based on models of STX binding to the channel pore and structure-activity studies of toxin binding^35,36^, we speculated that attachment of a sterically large, anionic photo-protecting group to STX-ea **1** would significantly destabilize toxin binding. The outer mouth of the channel funnels to the selectivity filter (the narrowest region of the ion permeation pathway), and STX is tightly lodged in this receptor site^35^. In addition, multiple aspartate and glutamate groups surround the toxin. Accordingly, conjugation of two different anionically-charged coumarin protecting groups, 6-bromo-7-hydroxy-4-(hydroxymethyl)coumarin (pK_a_ = 6.2, λ_max_ ~375 nm) ^37^, ^38^, ^39^ and 7-bis(carboxymethyl)-4-(hydroxymethyl)coumarin (λ_max_ ~380 nm)^33, 40^, to **1** were prioritized, yielding compounds **3** and **5**, respectively. Two additional coumarin derivatives, **2**and **4**, were prepared for comparative purposes and to better understand the requisite structural modifications that destabilize binding of the photocaged toxin to Na ^41,42^.

### Validation of photocaged STX derivatives

All photo-protected STXs were evaluated for binding affinity by whole-cell electrophysiology against Chinese hamster ovary (CHO) cells stably expressing rNaV1.2. Compounds **2**–**5** were up to 70-times less potent than STX-ea **1**, with **5** displaying the largest IC_50_ value of >1 μM (**Figure 2A**). As predicted, photocaged STXs bearing anionic groups are less potent than their uncharged counterparts (IC_50_: **2** = 27 nM vs **4** = 56 nM; **3** = 308 nM vs **5** = 1.004 μM). Similarly, increasing steric bulk also impairs toxin binding (*cf*. **4** and **5** vs **2** and **3**).

**Fig. 2:**
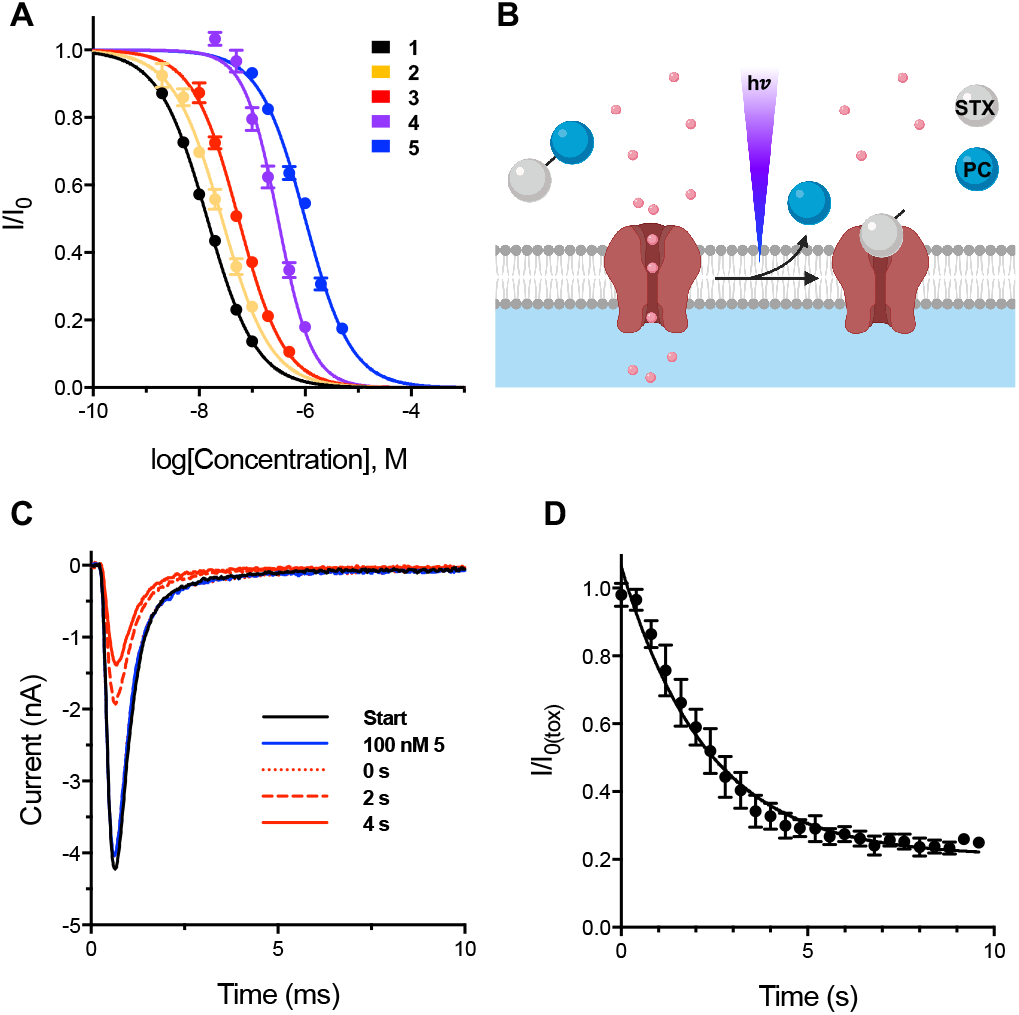
Photochemical release of STX-ea **1** results in block of NaV1.2 CHO. (**A**) Electrophysiological characterization of photocaged STXs against NaV1.2 CHO. IC_50_s, Hill Coefficients: **1** = 14.4 ± 0.3 nM, –0.94 ± 0.02; **2**= 27.0 ± 1.4 nM, –0.94 ± 0.05 (n = 3); **3** = 55.5 ± 2.1 nM, –1.01 ± 0.04; **4**= 307.5 ± 16.6 nM, –1.37 ± 0.09; **5** = 1003.8 ± 42.2 nM, –1.00 ± 0.04. Data represent mean ± s.e.m. n ≥ 5 unless otherwise noted. (**B**) Schematic depicting photolysis reaction to release **1** and subsequent channel block. Created with BioRender.com. (**C**) Representative trace depicting uncaging of 100 nM **5** against NaV1.2 CHO. Traces collected in the order: Start, 100 nM **5**, 0 s, 2 s, 4 s. Laser applied immediately prior to 10 ms, 0 mV voltage step (trace 0 s). (**D**) Time course of uncaging of 100 nM **5** against NaV1.2 CHO pulsed from –100 mV to 0 mV at 2.5 Hz. Laser was applied at t = 0 s. Data was subjected to exponential regression yielding τ = 2.3 ± 0.2 seconds (half-life 1.6 ± 0.1 seconds), R^2^ = 0.9056. (n = 4, mean ± s.e.m.).

In order to evaluate the uncaging efficiencies of photocaged STXs, voltage-clamped NaV1.2 CHO cells bathed in reagent were pulsed with 355 nm light (5 × 5 ms) while monitoring changes in channel block (**Figure 2B**). STX-eac **5** exhibited the highest uncaging efficiency, as measured by comparing the change in IC_50_ of the caged compound against the apparent IC_50_ following light exposure. For **5**, 5 × 5 ms of light exposure was highly effective at removing the coumarin group (**5** = 1.004 μM vs. 11 nM post-uncaging). By comparison, compounds **2**, **3**, and **4** displayed ≤ 3-fold change in IC_50_ under these same conditions (**Extended Data Figure 2**). Control experiments with NaV1.2 subjected to either laser light or STX-ea **1** showed no changes in sodium channel activity or peak current (**Extended Data Figure 1A–C**).

Following light exposure, all coumarin-modified STXs uncage and achieve maximal block of NaV1.2 within approximately four seconds (**Figure 2C**). Time course experiments with the most promising photocaged toxin, **5**, revealed a time constant for exponential current decay of 2.3 ± 0.2 seconds (i.e., half-maximal block achieved within 1.6 ± 0.1 seconds) (**Figure 2D**). The rapid time constant for block by the uncaged product, STX-ea **1**, is consistent with the measured association constant of this inhibitor (kon = 11.0 ± 0.7 × 10^6^ M^−1^s^−1^), which is similar to STX itself (**Extended Data Figure 1D-E**). Like STX, block of NaVs by **1** is fully reversible, as the toxin rapidly dissociates upon perfusion of cells with buffer solution (k_off_ = 3.8 × 10^2^ s^−1^).

### Interruption of action potential (AP) trains in primary neurons

To assess the utility of photocaged STX-ea to block APs, STX-eac **5** was evaluated against cultured rat embryonic hippocampal neurons (E18). In order to preserve cell health, laser excitation at 355 nm was limited to a 1 × 5 ms pulse. Against NaVs in primary neurons, the IC_50_ value for **5** was determined to be 2.148 ± 0.210 μM. The 2-fold larger IC_50_ value compared to that obtained against NaV1.2 in CHO cells may be due to the presence of multiple STX-sensitive NaV isoforms (1.1–1.3, 1.6)^43^ against which the potency of **5** is slightly varied (**Extended Data Figure 3**) or from the co-expression of β-auxiliary proteins (and/or other accessory proteins) in hippocampal neurons^35^. Analogous to experiments using CHO cells, the uncaging efficiency of **5** is high, as the apparent IC_50_ drops from 2.1 μM to 106 nM following a single 5 ms laser pulse (**Figure 3A-B**). Maximum channel inhibition is achieved after two seconds with a time constant of approximately one second (**Extended Data Figure 4A**) and is fully reversible upon toxin wash-off.

**Figure 3:**
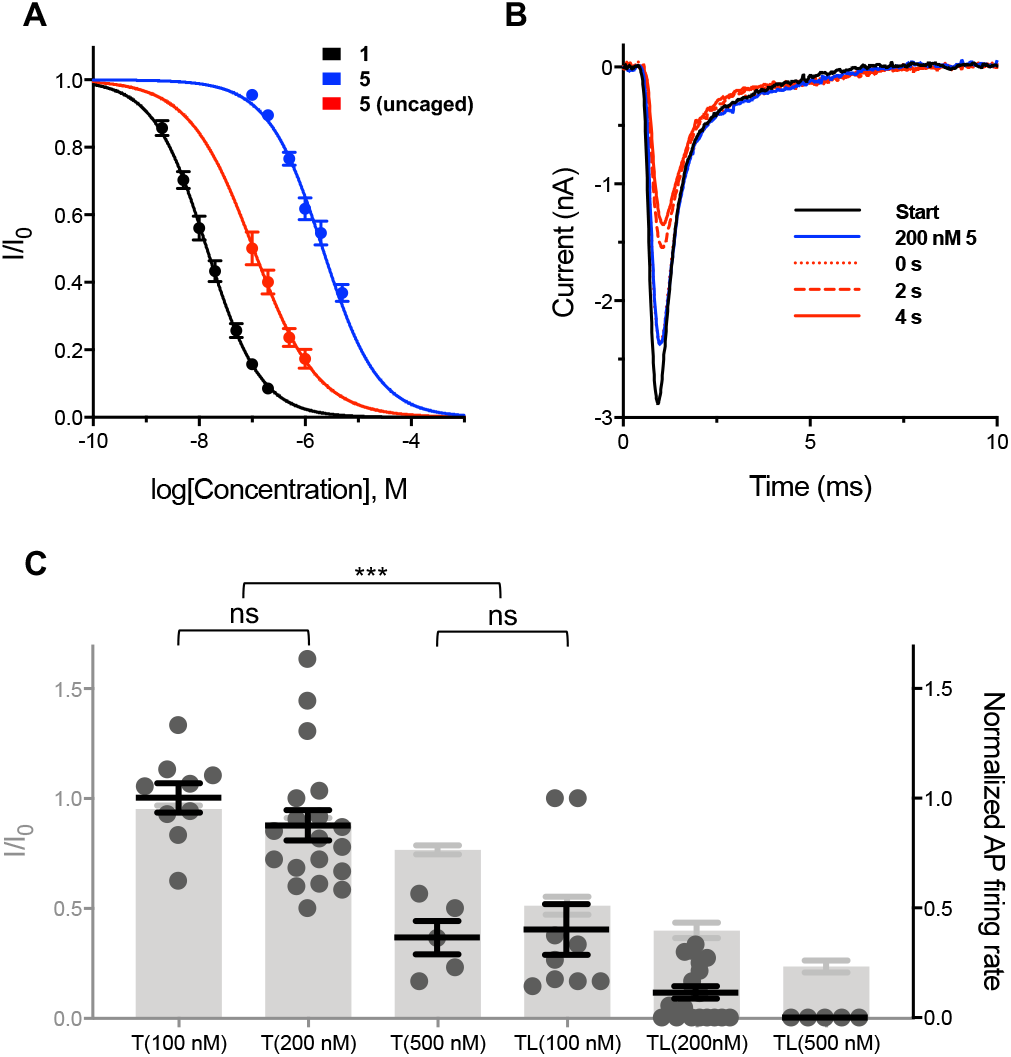
Uncaging of STX-eac **5** results in fast, concentration-dependent NaV block and inhibition of APs. (**A**) Electrophysiological characterization of photocaged STXs against hippocampal neurons DIV 6–8. IC_50_s, Hill Coefficients: **1**= 14.1 ± 0.8 nM, –0.86 ± 0.04; STX-eac **5** = 2148.4 ± 209.9 nM, –0.83 ± 0.07. Apparent IC_50_, Hill Coefficient: **5** (**uncaged**) = 106.0 ± 17.4 nM, –0.71 ± 0.11. Data represent mean ± s.e.m. n ≥ 5 unless otherwise noted. (**B**) Representative trace depicting uncaging of 200 nM **5** against hippocampal neurons DIV 6–8. Traces collected in the order: Start, 200 nM **5**, 0 s, 2 s, 4 s. Laser applied immediately prior to 10 ms, 0 mV voltage step (trace 0 s). (**C**) Comparison between current-clamp (DIV 9–13, right axis, black) and voltage-clamp (DIV 6–8, left axis, grey) data collected pre- and post-uncaging at various concentrations of **5**. T, toxin applied; TL, toxin and laser applied. Statistics calculated for current clamp data (n > 5, ****P<0.0001, one-way ANOVA with Tukey’s correction, mean ± s.e.m.).

Given the effectiveness of **5** for inhibiting NaV activity in hippocampal neurons, we next tested the ability of this reagent to impede action potential (AP) trains. Current-clamped dissociated neurons were bathed in varying concentrations of **5** to monitor the effects on AP firing frequency (**Extended Data Figures 5-7**). Notably, at 100–200 nM concentration, **5** has no influence on AP firing rate, despite blocking ~5–10% of the total population of NaVs (**Figure 3C**). Subsequent uncaging, particularly in experiments using 200 nM **5**, dramatically reduced spike frequency, on a time-scale consistent with voltage-clamp recordings (τ ~1.1 seconds; **Extended Data Figure 4B**). Analysis of both the voltage- and current-clamp data from hippocampal neurons indicates that ≥75% channel block is needed to completely eliminate AP trains (**Figure 3C**). Notably, blocking ~10–25% of NaVs resulted in the steepest decline in AP firing rate. Thus, it is possible to vary the concentration of **5**, laser power, and/or exposure time to alter AP frequency. For example, 200 nM **5** subjected to a single 5 ms laser pulse produced a six-fold increase in NaV block—from 10% to 60% inhibition pre-to post-uncaging—the difference between complete maintenance of AP firing and reduction to 10.5 ± 2.9% of initial frequency, with over 40% of cells incapable of initiating a single action potential (**Figure 4A-B**). AP block extends upward of 15 seconds, and is completely reversible upon perfusing cells with either buffer solution or a solution of **5** (**Figure 4C**). These data demonstrate that uncaging of **5** functions as a molecular circuit breaker to rapidly block trains of APs.

**Figure 4:**
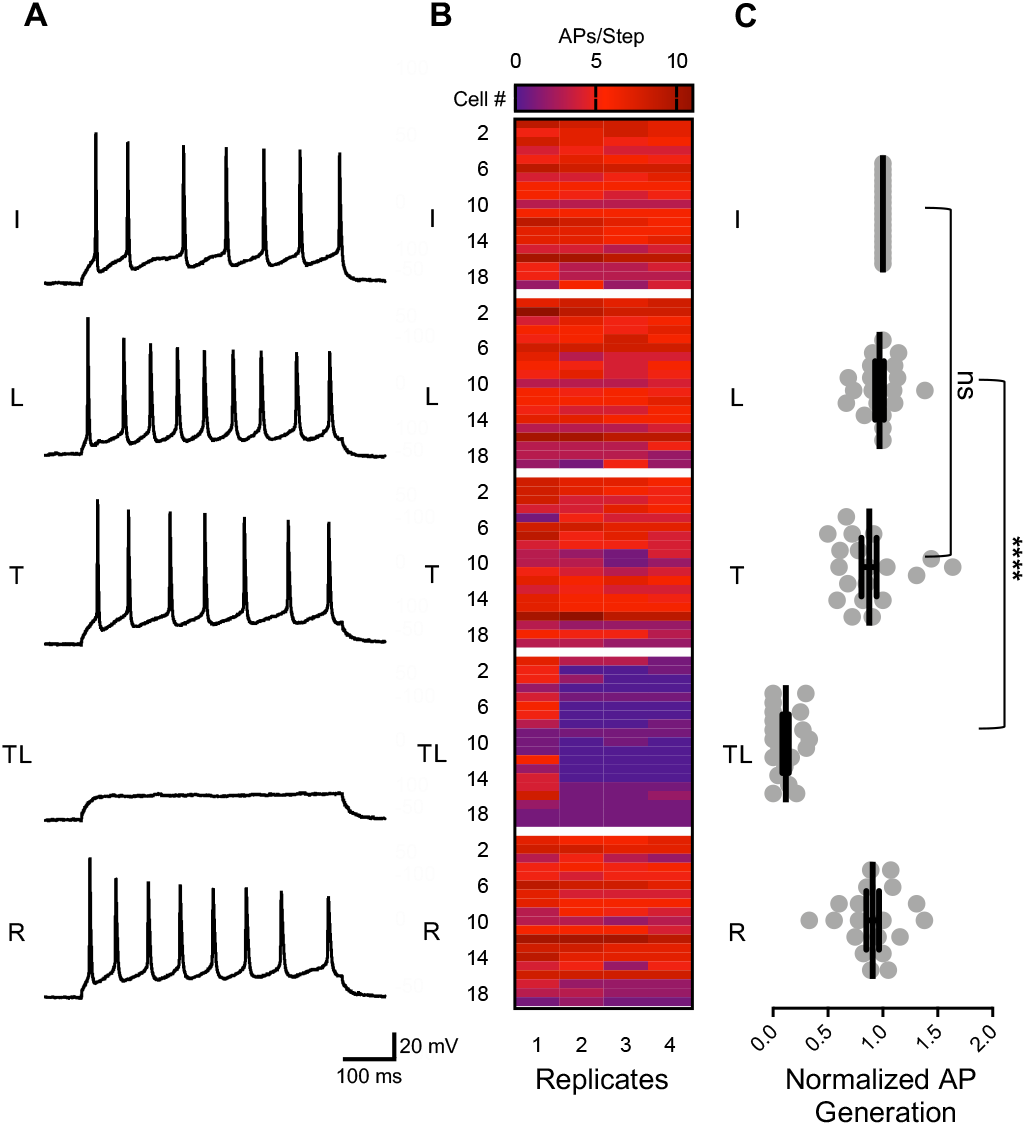
Uncaging of 200 nM STX-eac **5** shows reversible inhibition of AP firing. (**A**) Representative traces depicting initial (I), laser applied (L), 200 nM toxin **5** applied (T), 200 nM toxin **5** and laser applied (TL), and recovered (R) after wash-off I AP trains evoked by 500 ms, 50–150 pA current injections into hippocampal neurons DIV 9–13. Data taken from replicate current step 2 vis-à-vis **4B**. (**B**) Heat map summary of data described in (**A**) color-coded by number of action potentials per step (four replicate current steps at 0.25 Hz per condition, n = 19). (**C**) Equilibrated normalized action potential firing rate (i.e. over current steps 2–4) pre- and post-laser induced uncaging of **5** (n = 19, ****P<0.0001, one-way ANOVA with Tukey’s correction, mean ± s.e.m.).

### Effect of STX-eac in a corpus callosum slice preparation

To evaluate the functional effect of STX-eac **5** on axons, we performed electrophysiological analyses of compound action potentials (CAPs) evoked in the corpus callosum (**Figure 5A**). CAPs have been previously characterized in the mouse corpus callosum as a biphasic response with an early component (1-to 2-ms latency) representing mainly fast and large myelinated axons (N1) and a later component (3-to 6-ms latency) representing slower unmyelinated axons (N2)^44,45,46^ (**Figure 5B**). Previous studies of callosal signaling have been conducted at room temperature to slow down transmission speed in order to improve N1 detection^46^; our experiments were performed at 34 °C to more closely match physiological conditions. To improve detection of N1 and N2, we estimated current source density (CSD, **see methods**), which represents the inflow of Na^+^ into axons during the compound action potential. The first and second anti-peaks of the extracted CSD were used to determine the amplitude and timing of N1 and N2 action potential signals. This method of analysis was more accurate than measuring peak fiber volley in the local field potential (LFP) itself, as the latter was partially obscured by the electrical stimulus artifact. Examples of LFP and corresponding CSD traces are shown for 250 nM and 500 nM concentrations of **5** (**Figure 5C**). The amplitudes and timing of N1 and N2 peaks were determined for channels closest to the stimulating electrode.

**Figure 5:**
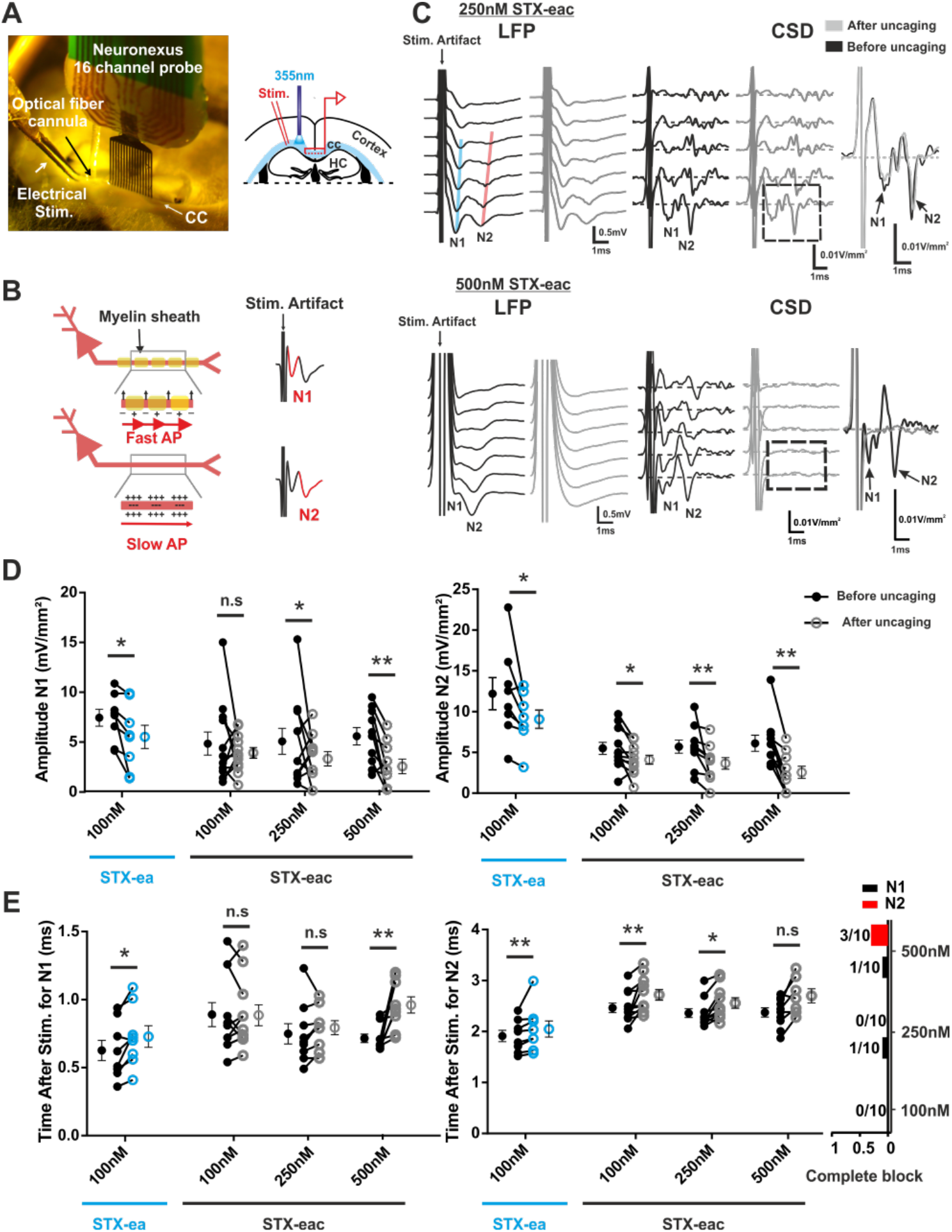
Uncaging STX-eac **5** differentially affects myelinated vs. unmyelinated callosal fiber transmission. (**A**) Experimental design of corpus callosum preparation. On the right, a representative picture of an experiment. On the left, a corresponding schematic representation^62^, HC stand for the hippocampus; cc for corpus callosum. (**B**) Schematic representation of callosal N1 and N2 features. N1 is the fastest component and corresponds mainly to myelinated axon fibers. N2 is slower and corresponds mainly to unmyelinated axon fibers. (**C**) Example of LFP and CSD signals for two different **5** concentrations, 250 nM (upper panel) and 500 nM (lower panel). Black traces represent signals before uncaging, grey traces represents signals after uncaging by 500 ms UV pulse light delivery. Only the first 7 channels closest to the stimulating electrode are represented. Expanded time scale of the first CSD channel is represented on the right. For 250 nM **5**, a reduction of amplitude and a delay of the peak time can be observed after uncaging for N2. Only a reduction of amplitude can be observed for N1; peak time was not affected. For 500 nM **5**, a loss of N1 and N2 signals can be observed after light delivery. (**D**) Scatter plot of the effect of 100 nM bath application of **1** in blue and of the effect of uncaging of 100 nM, 250 nM and 500 nM **5** on N1 (left graph) and N2 (right graph) amplitude. (**E**) Scatter plot of the effect 100 nM bath application of **1** in blue and of uncaging of 100 nM, 250 nM and 500 nM **5** on the N1 (left) and N2 (middle) peak time. Histograms (on the right) represent the incidence of complete block for N1 (black) and N2 (red) for the three different concentrations of **5**, 100 nM, 250 nM and 500 nM. Increases in peak timing were observed with 100 nM **1** and 500 nM **5** for N1 and with 100 nM **1**, 100 nM and 250 nM **5** for N2. Notice that 30 percent of the slices have a complete block of N2 at 500 nM, obscuring any potential difference in latency. (**D-E**) Wilcoxon, N=8, 7, 5 and 8 respectively for 100 nM **1**, 100 nM, 250 nM and 500 nM **5**. n.s. p > 0.05, *p < 0.05, **p < 0.01, ***.

In order to uncage **5**, an optical fiber was placed close to the stimulating electrode, in contact with the upper surface of the slice, to deliver UV light (**Figure 5A**). For each slice, we determined that exposure to UV light itself had a negligible effect on electrophysiological responses (**Extended Data Figure 8**). Quantitative analysis of peak amplitude and time revealed that the two CAP components, N1 and N2, are differentially affected by uncaging **5**. By design, **5** was focally uncaged along the path of the callosal fibers between the site of initiation in lateral corpus callosum and propagation was measured along the midline. Uncaging of 250 nM and 500 nM **5** produced N1 CAP amplitude decreases of 23% and 37%, respectively (**Extended Data Table 1**). N2 was more sensitive to uncaged **5**, with N2 peak sink amplitude reduced at all three concentrations of this compound––25% at 100 nM, 36% at 250 nM, and 58% at 500 nM (**Figure 5D**, **Extended Data Table 1**). These results indicate that transmission in unmyelinated callosal axons is more susceptible to partial sodium channel blockade than in myelinated fibers.

Our protocol for uncaging of **5** near the stimulation site revealed that callosal transmission speed (away from the uncaging location) along the midline is relatively insensitive to NaV block; however, the arrival time of the CAP at the recording electrode is delayed, especially for the N2 component. Peak latency for N1 was only affected after uncaging with the highest concentration of **5**, 500 nM, slowing by 0.28 ms. These results contrast rather starkly with recordings of N2, which show a delayed peak response of ~0.2 ms with 100 nM or 250 nM **5**, and suggest a slowed CAP conduction that is restricted to the portion of axons near the site of uncaging. The effect of uncaging 500 nM **5** on N2 CAP peak onset could not be reliably determined, as the signal is undetectable in 30% of these measurements (**Figures 5C, 5E**, **Extended Data Table 1**). Although signal arrival time of the CAP at the callosal midline was reduced by uncaged **5**, callosal fiber velocity, as estimated by the peak latency recorded along the midline of the callosum, was largely unaffected (**Extended Data Figure 8**, **Extended Data Table 1**).

Our findings demonstrate that focally uncaging **5** enables studies of action potential initiation and propagation, and that this tool compound could be particularly useful for modulating CAPs in unmyelinated axons. We wondered, however, if the differential effect of focally uncaged **5** on N1 vs. N2 might be due to an underlying sensitivity in the NaVs responsible for action potential initiation or propagation in myelinated vs unmyelinated fibers. To address this possibility, we tested the sensitivity of N1 and N2 to global, steady-state (i.e., bath) application of the active parent compound, STX-ea **1**. Quantitative analysis of peak amplitude (**Figure 5D**) and onset revealed (**Figure 5E**) that N1 and N2 are similarly affected by bath application of 100 nM **1**, which decreases N1 and N2 CAP amplitudes by 26% (**Extended Data Table 2**). In addition, bath application of **1** consistently affects callosal propagation speed, with a delayed peak response at the nearest electrode of both N1 and N2 CAP by approximately 0.1 ms each. In contrast with targeted sodium channel inhibition by photo-uncaging **5**, CAP velocity is significantly slowed for both N1 and N2 with bath application of **1**: from 1.36 ± 0.16 m/s to 1.00 ± 0.08 m/s for N1, and from 0.53 ± 0.01 m/s to 0.43 ± 0.03 m/s for N2 (**Extended Data Table 2**).

## Discussion

STX-eac **5** is the first molecular tool capable of spatiotemporal inhibition of NaVs to block the transmission of action potentials. Application of **5** to primary neuronal cells and tissue does not alter endogenous electrical signaling. Focal uncaging drives rapid release of STX-ea **1**, which achieves maximal NaV inhibition within two seconds following laser excitation. The percent of NaV block can be regulated by attenuating the laser power and duration of light exposure in addition to the concentration of **5**. We have demonstrated that **5** is well tolerated by cells for prolonged periods of time and that the effects of uncaging are fully reversible, with peak sodium currents restored within seconds to minutes upon perfusion of cells or tissue slices with buffer solution.

In cultured hippocampal neurons and in brain slice, **5** functions as a reversible molecular switch on AP generation. Due to the large potency difference between **5**(IC_50_ = 2.1 μM) and the uncaged compound, **1** (IC_50_ = 14 nM), complete block of evoked AP trains can be achieved following light exposure. As STX-sensitive NaV isoforms are preponderant in CNS tissue^47^, we anticipate that STX-eac **5** will be a valuable tool for examining NaV activity across a broad range of cell types. Moreover, given the choice of a coumarin photo-protecting group and the reported two-photon cross-sectional efficiency of uncaging such molecules (δ_unc_ > 1 GM at 740 nm^42^), we expect **5** to be particularly beneficial for studying subcellular populations of NaVs. While other coumarin-caged tool compounds (e.g., GABA-coumarin) have enjoyed extensive use in two-photon experiments^48^, **5** is the first such reagent capable of direct inhibition of action potential initiation. As such, STX-eac **5** offers a valuable compliment to photocaged neurotransmitters^49^ that have proven indispensable for investigations of the nervous system.

NaV blockade by uncaged STX-eac **5** is spatially precise. We have observed that UV uncaging is focal, as optical fiber placement must be within 300 μm of the callosum to differentiate the two components of the callosal CAP, N1 and N2. Focal uncaging of **5** at a precise location along the callosal tract reduces the amplitude and delays arrival of the N2 signal, but does not alter propagation speed in the region where the APs are measured (i.e., outside of the region of uncaging). Control experiments involving bath application of STX-ea **1** show that reduction in action potential amplitude *and speed* occurs in both N1 and N2 under these conditions. Accordingly, these results demonstrate that the effect of uncaging **5** is localized, as the measured AP propagation speed would be reduced if NaV block was not restricted to the site of light application.

Our data from corpus callosum recordings indicate that electrical transmission in unmyelinated fibers is more susceptible to attenuation than in myelinated fibers. We demonstrate that the faster conducting *in vitro* corpus callosum signal N1 is insensitive to localized delivery of STX-ea **1**, whereas the slower signal N2 from unmyelinated fiber is reduced in amplitude and timing. These findings have multiple potential explanations: First, AP conduction in unmyelinated fibers, which tend to be of smaller diameter^50^, may be more susceptible to local blockade. Modeling studies have suggested that NaV activity more significantly influences membrane voltage in thinner axon fibers (i.e., unmyelinated N2) than in larger axons (i.e., myelinated N1)^51^, thus local block of even a small fraction of NaVs could have a larger relative effect on small, unmyelinated fibers. A second contributing factor may stem from the different mechanisms of AP propagation in the two types of fibers. The corpus callosum contains diverse types and sizes of axons, divided into roughly two groups^52^. Larger diameter axons tend to be myelinated and more reliably transmit action potential due to saltatory conduction between insulated points (nodes of Ranvier), each expressing a high NaV density^53^. Inhibition of a small number of nodes can have a relatively minor effect on AP signaling, as saltatory propagation can apparently bypass or jump across short inactive segments^54^.

We have prepared STX-eac **5** and demonstrated the value of this high precision tool compound for localized block of NaVs in dissociated neurons and brain slice. When focally uncaged by a millisecond light pulse, release of the potent STX derivative, STX-ea **1**, results in rapid NaV inhibition and impedance of APs. In slice recordings on corpus collosum tissue, focal uncaging of **5** affords specific modulation of APs in slower-conducting, non-myelinated fibers. Future studies using this reagent may offer insight into the role of unmyelinated fibers in callosal signaling^50,55,56^ or the contributions of such fibers in demyelinating diseases such as multiple sclerosis, among other outstanding questions. We expect the availability of **5** to complement existing technologies for optical control of axonal signaling.

## Acknowledgements

This work was supported by a National Institutes of Health grant R21NS107003 to J.R.H and J.D.B and in part by R01 NS045684A1 (J.D.B). C.D.M was supported by the National Institutes of Health, National Institute of Neurological Disorders and Stroke K99NS104215. A.V.E. was supported by the Stanford Center for Molecular Analysis and Design (CMAD) as well as a Stanford Interdisciplinary Graduate Fellowship (SIGF) through the Stanford Bio-X Interdisciplinary Biosciences Institute. A.L.H. was supported by a Stanford Chemistry Undergraduate Summer Research Fellowship.

## Author Contributions

A.V.E, C.D.M, J.R.H., and J.D. conceived the project. A.V.E. and J.D. designed photocaged STXs. A.V.E. and A.L.H. synthesized all photocaged STXs and chemical intermediates. A.V.E. collected potency and uncaging data against NaV1.2 CHO cells, CHO-K1 cells, and singly dissociated hippocampal neurons. G.D. performed all slice physiology experiments. All authors contributed to data analysis and the writing of the manuscript.

## Methods

### Synthesis

#### General

All reagents were obtained commercially unless otherwise noted. Organic solutions were concentrated under reduced pressure (ca. 60 Torr) by rotary evaporation. Anhydrous CH2Cl2 and HPLC-grade CH3CN were obtained from commercial suppliers and used as is. *N,N*-Dimethylformamide (DMF) was passed through two columns of activated alumina prior to use. Triethylamine was distilled from calcium hydride.

Product purification was accomplished using forced-flow chromatography on Silicycle ultrapure silica gel (40–63 μm). Semi-preparative high-performance liquid chromatography (HPLC) was performed on a Varian ProStar model 210. Thin layer chromatography was performed on EM Science silica gel 60 F_254_ plates (250 μm). Visualization of the developed chromatogram was accomplished by fluorescence quenching. High-resolution mass spectra were obtained from the Vincent Coates Foundation Mass Spectrometry Laboratory at Stanford University. Samples were analyzed with LC/ESI-MS by direct injection onto a Waters Acquity UPLC and Thermo Fisher Exactive mass spectrometer scanning m/z 100– 2000. The LC mobile phase was 100% methanol and the flow rate was 0.175 mL/min.

Saxitoxin derivatives were quantified by ^1^H NMR spectroscopy on a Varian Inova 600 MHz NMR instrument using distilled DMF as an internal standard. A relaxation delay (d1) of 20 s and an acquisition time (at) of 10 s were used for spectral acquisition. The concentration of the toxin derivative was determined by integration of ^1^H signals corresponding to toxin and a fixed concentration of the DMF standard.

#### Compounds

Coumarins 6-bromo-4-(hydroxymethyl)-7-methoxycoumarin (**6**), 6-bromo-7-hydroxy-4-(hydroxymethyl)coumarin (**7**), and 7-[bis(tert-butoxycarbonylmethyl)-amino]-4-(hydroxymethyl)coumarin (**8**) were prepared as described in Furuta, *et al.*^41^, Furuta, *et al.*^37^, and Noguchi, *et al.*^57^, respectively.

Synthesis of N21-saxitoxin ethylamine (**1**) was adapted from our previously disclosed work^31^. N-Hydroxysuccinimide esters and photocaged STXs were synthesized as described in the extended data.

### Plasmids

Oocyte expression vector pLCT2-rNaV 1.3 was a generous gift from Dr. A. L. Goldin (University of California, Irvine, Department of Microbiology & Molecular Genetics). The full-length cDNA encoding for the alpha subunit of rNaV 1.3 was excised and inserted into a low-copy modified pcDNA3.1(+) vector^58^ by VectorBuilder (Vector ID VB190704-1006vdy). Detailed information about the vector can be retrieved at vectorbuilder.com. Mammalian expression vector pZem228 containing the cDNA coding for the alpha subunit of rat NaV 1.4 was a generous gift from Dr. S. R. Levinson (University of Colorado, Department of Physiology and Biophysics). Mammalian expression vector pcDNA3.1(+) containing the cDNA coding for the alpha subunit of human NaV1.5 originated from Dr. T. R. Cummins (Indiana University, Department of Biology).

### Cell Culture

#### Chinese hamster ovary (CHO) cells stably expressing rat NaV 1.2

Stably-expressing NaV1.2 CHO cells were generously provided by Dr. W. A. Catterall (University of Washington, Department of Pharmacology). Cells were grown on 10 cm tissue culture dishes in RPMI 1640 medium with L-glutamine (Thermo Fisher, Waltham, MA) and supplemented with 10% fetal bovine serum (ATCC, Manassas, VA), 50 U/mL penicillin-streptomycin (Thermo Fisher, Waltham, MA), and 0.2 μg/mL G418 (Sigma-Aldrich Co., St. Louis, MO). Cells were kept in a 37 °C, 5% carbon dioxide, 96% relative humidity incubator and passaged approximately every three days. Passaging of cells was accomplished by aspiration of media, washing with phosphate-buffered saline, treatment with 1 mL of trypsin-EDTA (0.05% trypsin, Millipore Sigma, Hayward, CA) until full dissociation of cells was observed (approximately five minutes), and dilution with 4 mL of growth medium. Cells were routinely passaged at 1 in 20 to 1 in 10 dilution.

#### Chinese hamster ovary K1 (CHO-K1) cells transiently expressing NaVs

CHO-K1 cells were grown on 10 cm tissue culture dishes in F12-K medium (ATCC, Manassas, VA) and supplemented with 10% fetal bovine serum (ATCC, Manassas, VA) and 50 U/mL penicillin-streptomycin (Thermo Fisher, Waltham, MA). Cells were kept in a 37 °C, 5% carbon dioxide, 96% relative humidity incubator and passaged approximately every three days. Passaging of cells was accomplished by aspiration of media, washing with phosphate-buffered saline, treatment with 1 mL of trypsin-EDTA (0.05% trypsin, Millipore Sigma, Hayward, CA) until full dissociation of cells was observed (approximately five minutes), and dilution with 4 mL of growth medium. Cells were routinely passaged at 1 in 20 to 1 in 10 dilution. Lipofectamine™ LTX PLUS™ Reagent was used to accomplish all transient transfections, according to the manufacturer’s instructions (Thermo Fisher, Waltham, MA).

#### Rat embryonic day 18 hippocampal neurons

Prior to dissection, 5 mm diameter, 0.15 mm thick round glass coverslips (Warner Instruments, Hamden, CT) were coated overnight with 1 mg/mL poly-D-lysine hydrobromide (PDL, molecular weight 70,000–150,000 Da, Millipore Sigma, Hayward, CA) in 0.1 M, pH 8.5 borate buffer in a 37 °C, 5% carbon dioxide, 96% relative humidity incubator.

Hippocampi were dissected from embryonic day 18 fetuses into ice-cold Hibernate E (BrainBits, LLC, Springfield, IL) as previously described^59^. Following dissection, cells were dissociated in 2 mL of trypsin-EDTA for 15 minutes in a 37 °C, 5% carbon dioxide, 96% relative humidity incubator. Subsequently, trypsinized cells were quenched with 10 mL of quenching medium (DMEM high glucose (Thermo Fisher, Waltham, MA) supplemented with 15% fetal bovine serum, 100 U/mL penicillin-streptomycin, and 1 mM MEM sodium pyruvate (Atlanta biologicals, Flowery Branch, GA)). Tissue was allowed to settle, supernatant was removed, and the tissue pellet was rinsed twice more with 10 mL of quenching medium. Cells were then manually dissociated into 2 mL of plating medium (DMEM supplemented with 10% FBS, 50 U/mL penicillin-streptomycin, and 1 mM MEM sodium pyruvate) by pipetting with a fire-polished 9” borosiligate glass Pasteur pipet (Fisher Scientific, Waltham, MA).

Cells were plated onto PDL-coated 5 mm glass coverslips in 35 mm tissue culture dishes containing 2 mL of plating medium at a density of 200,000 cells/dish (for voltage-clamp experiments) or 600,000 cells/dish (for current-clamp experiments). After 45 minutes, coverslips were transferred to new tissue culture dishes containing 2 mL of maintenance medium (neurobasal supplemented with 1x B-27 Supplement, 1x Glutamax, and 50 U/mL penicillin-streptomycin (Thermo Fisher, Waltham, MA)). Cells were fed every 3– 4 days by changing 50% of the working medium.

### Single cell electrophysiology

Data were measured using the patch-clamp technique in the whole-cell configuration with an Axon Axopatch 200B amplifier (Molecular Devices, San Jose, CA). The output of the patch-clamp amplifier was filtered with a built-in lowpass, four-pole Bessel filter having a cutoff frequency of 5 kHz for voltage-clamp recordings or 10 kHz for current-clamp recordings, and sampled at 100 kHz. Pulse stimulation and data acquisition used Molecular Devices Digidata 1322A or 1550B controlled with pCLAMP software version 10.4 or 11.1, respectively (Molecular Devices, San Jose, CA).

Borosilicate glass micropipettes (Sutter Instruments, Novato, CA) were fire-polished to a tip diameter yielding a resistance of 1.3–5.5 MΩ, for NaV1.2 CHO cells, or 3.0–9.0 MΩ, for E18 hippocampal neurons, in the working solutions.

#### Voltage-clamp recordings

For NaV1.2 CHO and CHO-K1 cells, the internal solution was composed of 40 mM NaF, 1 mM EDTA, 20 mM HEPES, and 125 mM CsCl (pH 7.4 with CsOH); the external solution comprised 160 mM NaCl, 2 mM CaCl2, and 20 mM HEPES (pH 7.4 with CsOH). For E18, DIV 6–8 hippocampal neurons, the internal solution was composed of 114.5 mM gluconic acid, 114.5 mM CsOH, 2 mM NaCl, 8 mM CsCl, 10 mM MOPS, 4 mM EGTA, 4 mM MgATP, and 0.3 mM Na2GTP (pH 7.3 with CsOH, 240 mOsm with glucose), while the external solution was Hibernate E low fluorescence (BrainBits, LLC, Springfield, IL).

Currents were elicited by 10 ms step depolarizations from a holding potential (–100 mV for NaV1.2 CHO and CHO-K1 cells expressing NaV1.4, –120 mV for CHO-K1 cells expressing NaV1.3 or 1.5, or –80 mV for E18 hippocampal neurons) to 0 mV at a rate of 0.5 Hz (unless otherwise noted). Leak currents were subtracted using a standard P/4 protocol of the same polarity. Series resistance was compensated at 80–95% with a τ_lag_ of 20 or 35 ms for NaV1.2 CHO and CHO-K1 cells or E18 hippocampal neurons, respectively. All measurements were recorded at room temperature (20–25 °C). Data were normalized to control currents, plotted against toxin concentration, and analyzed using Prism 8 (GraphPad Software, LLC). Data were fit to concentration-response curves to obtain IC_50_ values and expressed as mean ± s.e.m. The number of observations (n) was ≥ 5 for all reported data unless otherwise noted.

#### Current-clamp recordings

Data were collected on DIV 9–13 hippocampal neurons firing action potential trains with frequencies greater than 5 Hz. The internal solution was composed of 130 mM CH_3_SO_3_–K^+^, 8 mM NaCl, 10 mM HEPES, 10 mM Na_2_ phosphocreatine, 3 mM L-ascorbic acid, 4 mM MgATP, and 0.4 mM Na_2_GTP (pH 7.4 with KOH, 305 mOsm with glucose); the external solution comprised Hibernate E low fluorescence adjusted to 310 mOsm with 40 mM NaCl.

Action potentials were elicited by four 500 ms current injections of 50–150 pA at a rate of 0.25 Hz. Series resistance was typically compensated at 90–95% with a τ_lag_ of 35 ms. All measurements were recorded at room temperature (20–25 °C). Data were analyzed using Clampfit (Molecular Devices, San Jose, CA). The number of observations (n) was ≥ 5 for all reported data. Statistical comparisons were performed using two-tailed Student’s *t*-tests assuming unequal variances.

#### UV Laser (355 nm)

A pulsed 355 nm UV laser beam (DPSS Lasers, Model 3501-100) was directed through a 200 μm core optical fiber to a 200 μm core, 0.22 NA fiber optic cannula (Thorlabs, Newton, NJ) to the clamped cell. Unless otherwise noted, photolysis was induced by 5 ms, 130 mW UV pulses activated immediately prior to the depolarization (or current injection) step.

### Extracellular multielectrode recordings

Extracellular recordings were obtained with neocortical coronal 400 μm thick slices obtained from young (~P35) wild type mice. Slices containing the midline crossing segments of the corpus callosum (Bregma 0.4 to −1) were saved, which represents 3 slices per brain. Recordings were performed in a humidified oxygenated interface chamber at 34°C and perfused at rate of 2 ml/min with oxygenated ACSF. The ACSF contained: 10 mM glucose, 26 mM NaHCO3, 2.5 mM KCl, 1.25 mM NaHPO_4_, 1 mM MgSO_4_, 2 mM CaCl_2_, and 126 mM NaCl (298 mOsm).

A linear silicon multichannel probe (16 channels, 100 μm inter-electrode spacing, NeuroNexus Technologies) was placed in the midline of the corpus callosum. A bipolar tungsten microelectrode (each wire, 50-100 kΩ, FHC) was positioned in the corpus callosum in one hemisphere at approximatively 1 mm from the closest channel of the recording electrode, angled to ensure that both contacts were within the boundaries of the callosum. Both electrodes were initially lowered to approximately 200 μm slice depth, and fine adjustments were made to optimize the signal amplitude of the evoked compound action potential (CAP), representing the summed activity of callosal fibers^44^. Biphasic electrical stimuli (± 400 μA, 0.2 ms each phase) were delivered every 10 seconds to elicit the callosal CAP response. This evoked response has previously been characterized by a faster component generated mainly by fast-conducting myelinated axons (N1), and a slower component reflecting mainly slower unmyelinated axons (N2)^60^.

A laser pulse of 500 ms, 40 mW, 355 nm UV (DPSS Lasers, Model 3501-100) was directed through a 200 μm core optical fiber to a 0.22 NA cannula (Thorlabs, Newton, NJ) placed along the callosum between the recording and stimulating electrode to photo-uncage **5**. Three concentrations of **5**(100 nM, 250 nM and 500 nM) were tested on callosal CAP N1 and N2 in three different mice groups (respectively N=7, N=5 and N=8).

The following protocol was performed identically on each slice: Application of aCSF for 5 min, delivery of a UV light pulse, 5 min of aCSF, application of STX-eac for 10 min, delivery of a UV light pulse, 5 min of uncaged version of STX-eac, 10 min wash-off with aCSF.

For the bath application experiment, 100 nM **1**was used. After 5 min of aCSF application, callosal responses were recorded to be use as baseline. Then, **1**was applied in bath for 10 min and 15 callosal responses were elicited and recorded. Wash-off with aCSF was next performed during 10 min.

Evoked callosal CAP field potentials elicited by electrical stimulations were digitized at 25 kHz, and stored using RZ5D processor multichannel workstation (Tucker-Davis Technologies). Signals were band-pass filtered between 1 Hz and 3 kHz. To obtain a more reliable index of the location, direction, and magnitude of currents underlying synchronous network activity, we derived the current-source density (CSD) from raw LFP signals^61^. Assuming a uniform extracellular resistivity, the CSD can be estimated as is the second spatial derivative of the LFP. CSD peak amplitudes and times of N_1_ and N_2_ components of the closest channel to the stimulating electrode were quantified using trial-averaged responses (10 trials per condition).

