## Supplemental Information for "Precise spatiotemporal control of voltage-gated sodium channels by photocaged saxitoxin"

### Extended Data

1. Extended Data Figures
2. Extended Data Tables
3. Chemical Procedures
4. $^{1}$H NMR Spectra

#### 1. Extended Data Figures

**Extended Data Figure 1.** Laser application does not alter the $IC_{50}$ of **1** or peak current produced by $Na_V$s.

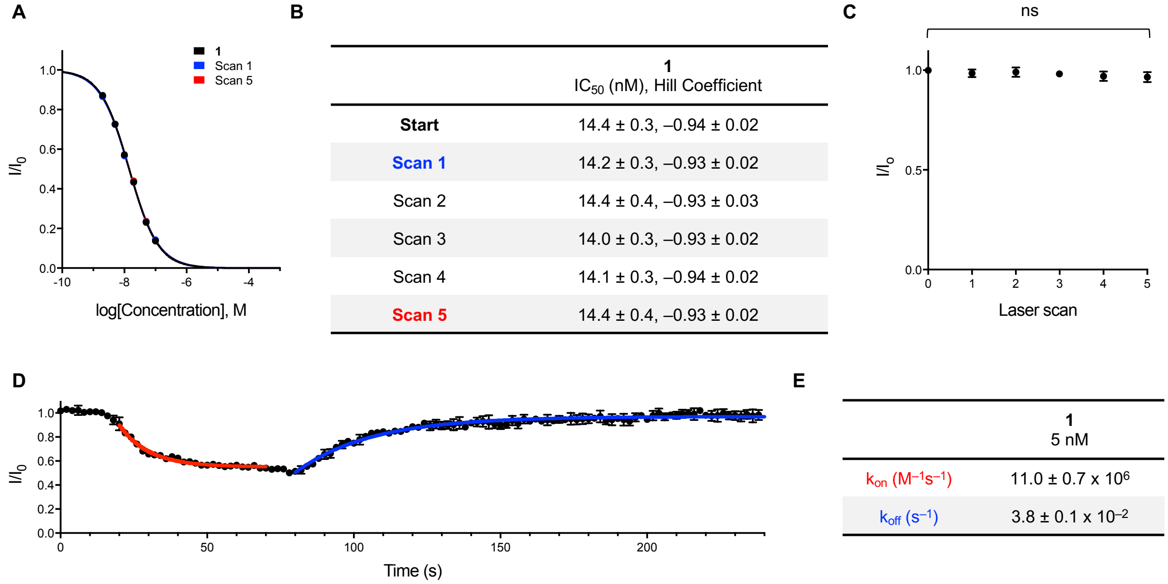


**B**

| | **1**<br>$IC_{50}$ (nM), Hill Coefficient |
|---|---|
| **Start** | 14.4 ± 0.3, –0.94 ± 0.02 |
| **Scan 1** | 14.2 ± 0.3, –0.93 ± 0.02 |
| Scan 2 | 14.4 ± 0.4, –0.93 ± 0.03 |
| Scan 3 | 14.0 ± 0.3, –0.93 ± 0.02 |
| Scan 4 | 14.1 ± 0.3, –0.94 ± 0.02 |
| **Scan 5** | 14.4 ± 0.4, –0.93 ± 0.02 |

**E**

| | **1**<br>5 nM |
|---|---|
| $k_{on}$ ($M^{-1}s^{-1}$) | 11.0 ± 0.7 x $10^6$ |
| $k_{off}$ ($s^{-1}$) | 3.8 ± 0.1 x $10^{-2}$ |

(**A**) Electrophysiological characterization of STX-ea **1** against $Na_V$1.2 CHO. Initial $IC_{50}$ in black, following laser scan 1 in blue, and after laser scan 5 in red. (**B**) Table showing apparent $IC_{50}$s of **1** pre- and post-laser photolysis for up to five laser scans. (**C**) Normalized total current produced by $Na_V$1.2 pre- and post-application of up to five laser scans (n = 7, one-way ANOVA with Tukey's correction, mean ± s.e.m.). (**D**) Time course of **1** binding to and wash-off from $Na_V$1.2 CHO. Toxin **1** (5 nM) was applied for one minute and washed out for three minutes at a constant perfusion rate of 1 mL/min. Channels were pulsed at 2.5 Hz (abridged data shown here). Data represent mean ± s.e.m. (n = 3). Binding data (t = 20–70 s) and wash-off data (t = 80–240 s) were subjected to independent exponential regressions (red and blue, respectively), yielding $\tau$(on) = 10.8 ± 0.4 s and $\tau$(off) = 26.4 ± 0.8 s. (**E**) Association and dissociation constants were calculated for **1** against $Na_V$1.2 CHO according to the equations by Hahn and Strichartz.[1]

**Extended Data Figure 2.** Photocaged STXs **3** and **5** uncage more effectively than **2** and **4**.

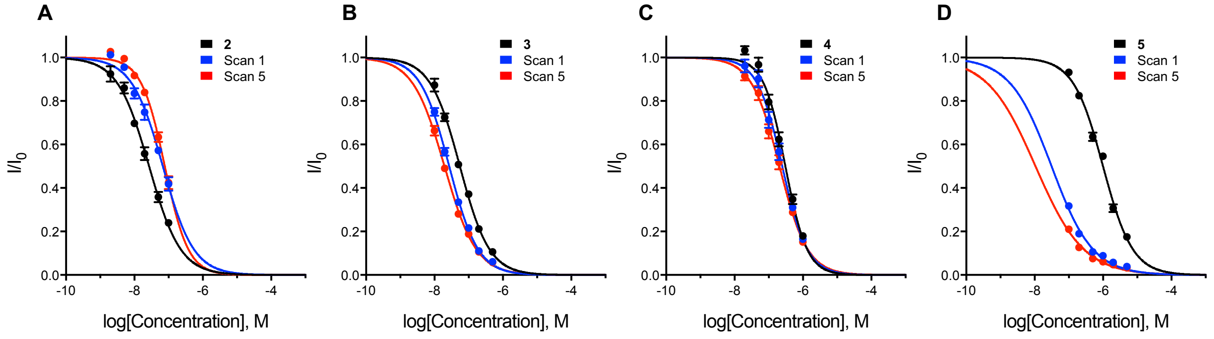


E

| | **2**<br>$IC_{50}$ (nM), Hill Coefficient | **3**<br>$IC_{50}$ (nM), Hill Coefficient | **4**<br>$IC_{50}$ (nM), Hill Coefficient | **5**<br>$IC_{50}$ (nM), Hill Coefficient |
|---|---|---|---|---|
| **Start** | 27.0 ± 1.4, –0.94 ± 0.05 | 55.5 ± 2.1, –1.01 ± 0.04 | 307.5 ± 16.6, –1.37 ± 0.09 | 1003.8 ± 42.2, –1.00 ± 0.04 |
| **Scan 1** | 68.1 ± 5.0, –0.96 ± 0.07 | 27.0 ± 0.8, –1.03 ± 0.03 | 248.9 ± 15.2, –1.19 ± 0.08 | 31.4 ± 3.3, –0.72 ± 0.04 |
| Scan 2 | 77.1 ± 5.3, –1.24 ± 0.12 | 22.3 ± 0.9, –0.96 ± 0.04 | 224.1 ± 12.7, –1.12 ± 0.07 | 13.4 ± 2.3, –0.62 ± 0.04 |
| Scan 3 | 78.6 ± 5.0, –1.31 ± 0.12 | 20.4 ± 0.7, –0.93 ± 0.03 | 223.8 ± 12.5, –1.08 ± 0.06 | 10.4 ± 1.8, –0.60 ± 0.04 |
| Scan 4 | 86.8 ± 5.6, –1.18 ± 0.10 | 19.8 ± 0.6, –0.91 ± 0.03 | 218.9 ± 11.2, –1.09 ± 0.06 | 11.2 ± 1.7, –0.62 ± 0.03 |
| **Scan 5** | 77.7 ± 3.8, –1.26 ± 0.08 | 19.8 ± 0.7, –0.94 ± 0.03 | 206.9 ± 11.0, –1.06 ± 0.06 | 10.7 ± 1.9, –0.62 ± 0.04 |

(**A**)–(**D**) Electrophysiological characterization of photocaged STXs **2**–**5** against $Na_V1.2$ CHO. Initial $IC_{50}$ in black; apparent $IC_{50}$ following laser scan 1 in blue and after laser scan 5 in red. (**E**) Table showing apparent $IC_{50}$s of photocaged STXs pre- and post-laser induced uncaging for up to five laser scans. Data represent mean ± s.e.m. (n = 3–6).

**Extended Data Figure 3.** Compounds **1** and **5** exhibit similar potencies against other STX-sensitive $Na_V$ isoforms.

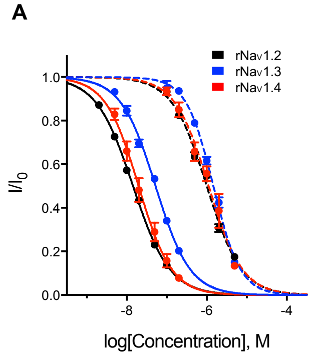


B

| | **1**<br>$IC_{50}$ (nM), Hill Coefficient | **5**<br>$IC_{50}$ (nM), Hill Coefficient |
|---|---|---|
| $rNa_V1.2$ | 14.4 ± 0.3, –0.94 ± 0.02 | 1003.8 ± 42.2, –1.00 ± 0.04 |
| $rNa_V1.3$ | 52.6 ± 1.9, –1.01 ± 0.04 | 1467.1 ± 54.5, –1.28 ± 0.06 |
| $rNa_V1.4$ | 19.8 ± 1.5, –1.04 ± 0.08 | 1120.4 ± 99.3, –1.03 ± 0.10 |
| $hNa_V1.5$ | >2 µM | >2 µM |

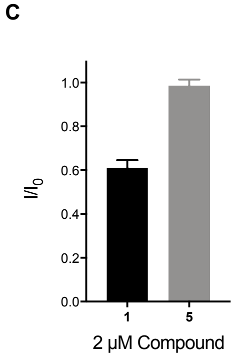


(**A**) Electrophysiological characterization of **1** (solid) and **5** (dashed) against $Na_V1.2$ CHO, CHO-K1 expressing $rNa_V1.3$, and CHO-K1 expressing $rNa_V1.4$. (**B**) Table showing $IC_{50}$s of **1** and **5**. (**C**) Electrophysiological characterization of **1** and **5** against CHO-K1 expressing $hNa_V1.5$; data collected at 2 µM. Data represent mean ± s.e.m. (n = 3–7).

**Extended Data Figure 4.** Uncaging of STX-eac **5** is functionally complete after four seconds against hippocampal neurons.

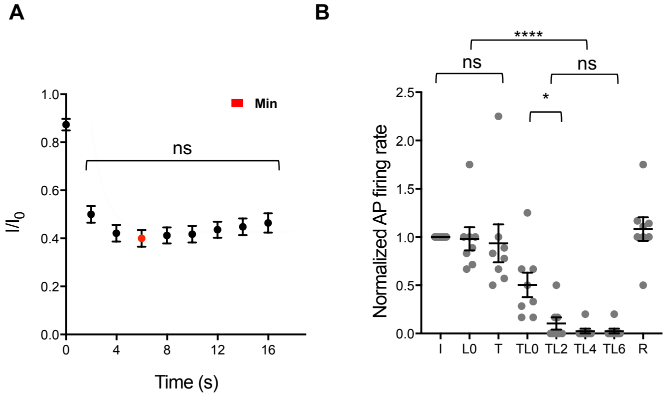


(**A**) Time course of uncaging of 200 nM **5** against hippocampal neurons DIV 6–8. Laser was applied at t = 0 s. Data was subjected to exponential regression yielding $\tau = 1.0 \pm 0.3$ seconds, $R^2 = 0.7521$. (n > 5, two-way ANOVA with Tukey's correction, mean ± s.e.m.). Data from Min used to plot **Figure 3**. (**B**) Time course of uncaging of 200 nM **5** against hippocampal neurons DIV 9–13. Graph depicts normalized AP firing rates evoked by 500 ms, 50–150 pA current injections either: initially (I), 0 seconds after laser application (L0), post-toxin application (T), post-toxin application and 0, 2, 4, or 6 seconds after laser application (TL0, TL2, TL4, and TL6, respectively), and post-wash off/recovery (R). Data was subjected to exponential regression yielding $\tau = 1.1 \pm 0.8$ seconds, $R^2 = 0.5124$. (n > 5, *P<0.05, ****P<0.0001, one-way ANOVA with Tukey's correction, mean ± s.e.m.).

**Extended Data Figure 5.** Uncaging of 100 nM STX-eac **5** with one 5 ms laser pulse is insufficient to consistently inhibit action potential trains in hippocampal neurons.

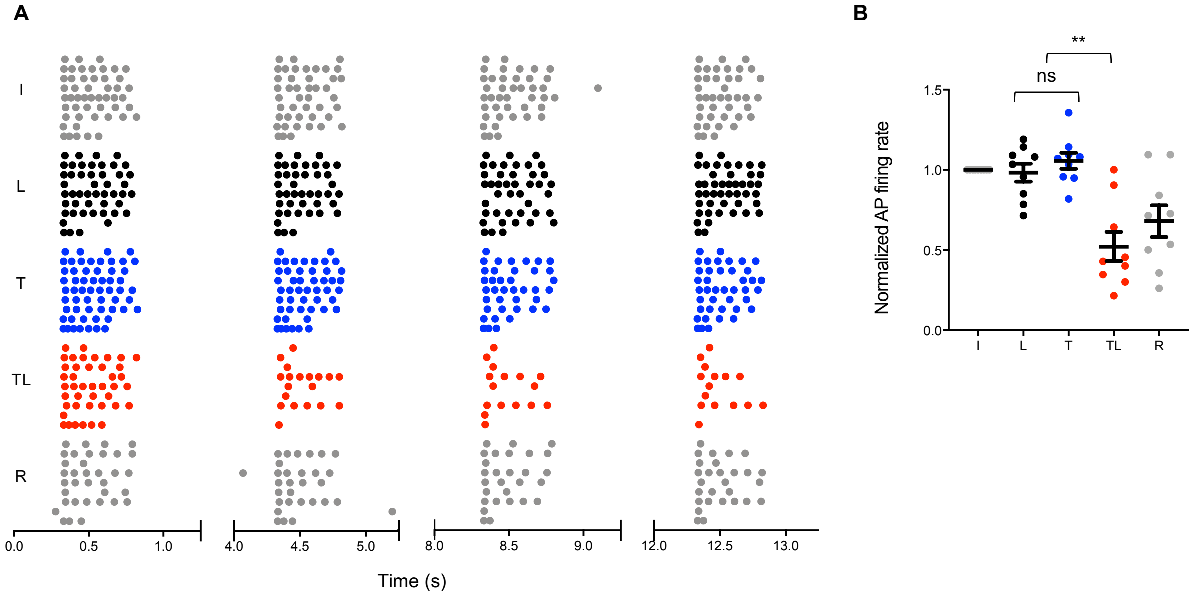


(**A**) Raster plot depicting initial (I), laser applied (L), 100 nM **5** applied (T), 100 nM STX-eac **5** and laser applied (TL), and recovered following wash off (R) action potential trains evoked by 500 ms, 50–150 pA, 0.25 Hz current injections into hippocampal neurons DIV 9–13. (**B**) Normalized action potential firing rates calculated for data in (**A**) (n = 9, **P<0.01, one-way ANOVA with Tukey's correction, mean ± s.e.m.).

**Extended Data Figure 6.** Uncaging of 200 nM STX-eac **5** with one 5 ms laser pulse is sufficient to consistently inhibit action potential trains in hippocampal neurons.

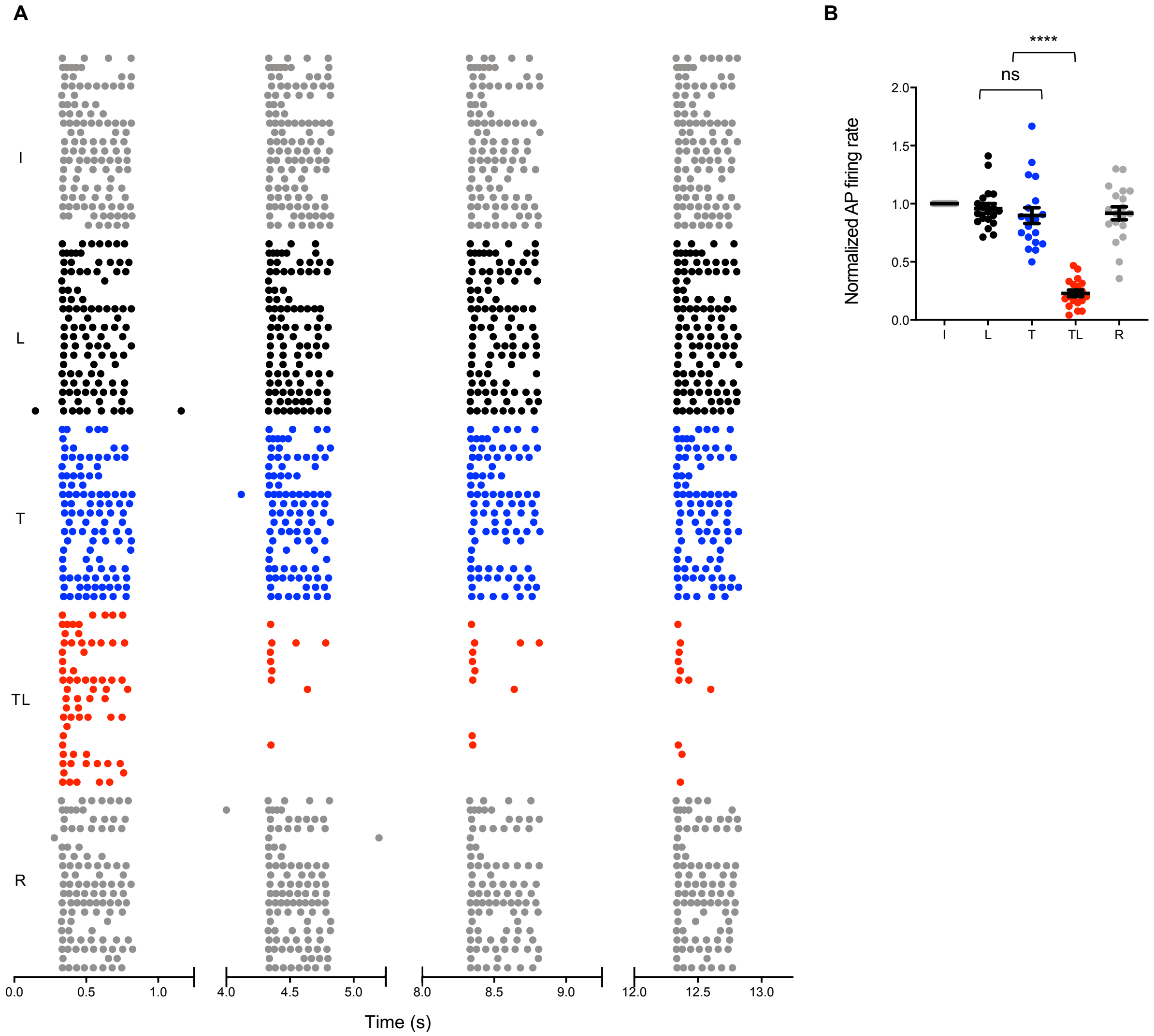


(**A**) Raster plot depicting initial (I), laser applied (L), 200 nM **5** applied (T), 200 nM STX-each **5** and laser applied (TL), and recovered following wash off (R) action potential trains evoked by 500 ms, 50–150 pA, 0.25 Hz current injections into hippocampal neurons DIV 9–13. (**B**) Normalized action potential firing rates calculated for data in (**A**) (n = 19, ****P<0.0001, one-way ANOVA with Tukey's correction, mean ± s.e.m.).

**Extended Data Figure 7.** 500 nM STX-eac **5** alters action potential trains absent laser-induced uncaging.

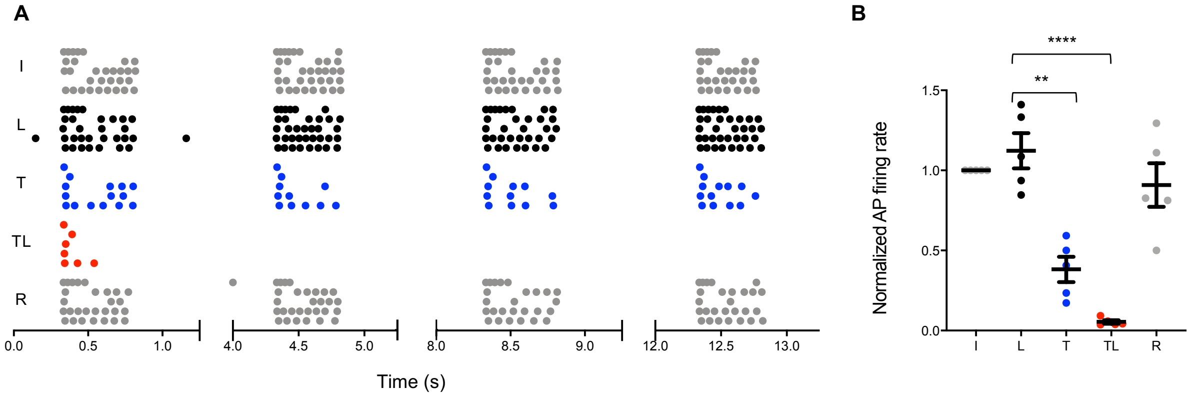


(**A**) Raster plot depicting initial (I), laser applied (L), 500 nM **5** applied (T), 500 nM STX-eac **5** and laser applied (TL), and recovered following wash off (R) action potential trains evoked by 500 ms, 50–150 pA, 0.25 Hz current injections into hippocampal neurons DIV 9–13. (**B**) Normalized action potential firing rates calculated for data in (**A**) (n = 5, **P<0.01, ****P<0.0001, one-way ANOVA with Tukey's correction, mean ± s.e.m.).

**Extended Data Figure 8.** Differential effect of 100 nM STX-ea **1** bath application and uncaged STX-eac **5** on callosal signal propagation.

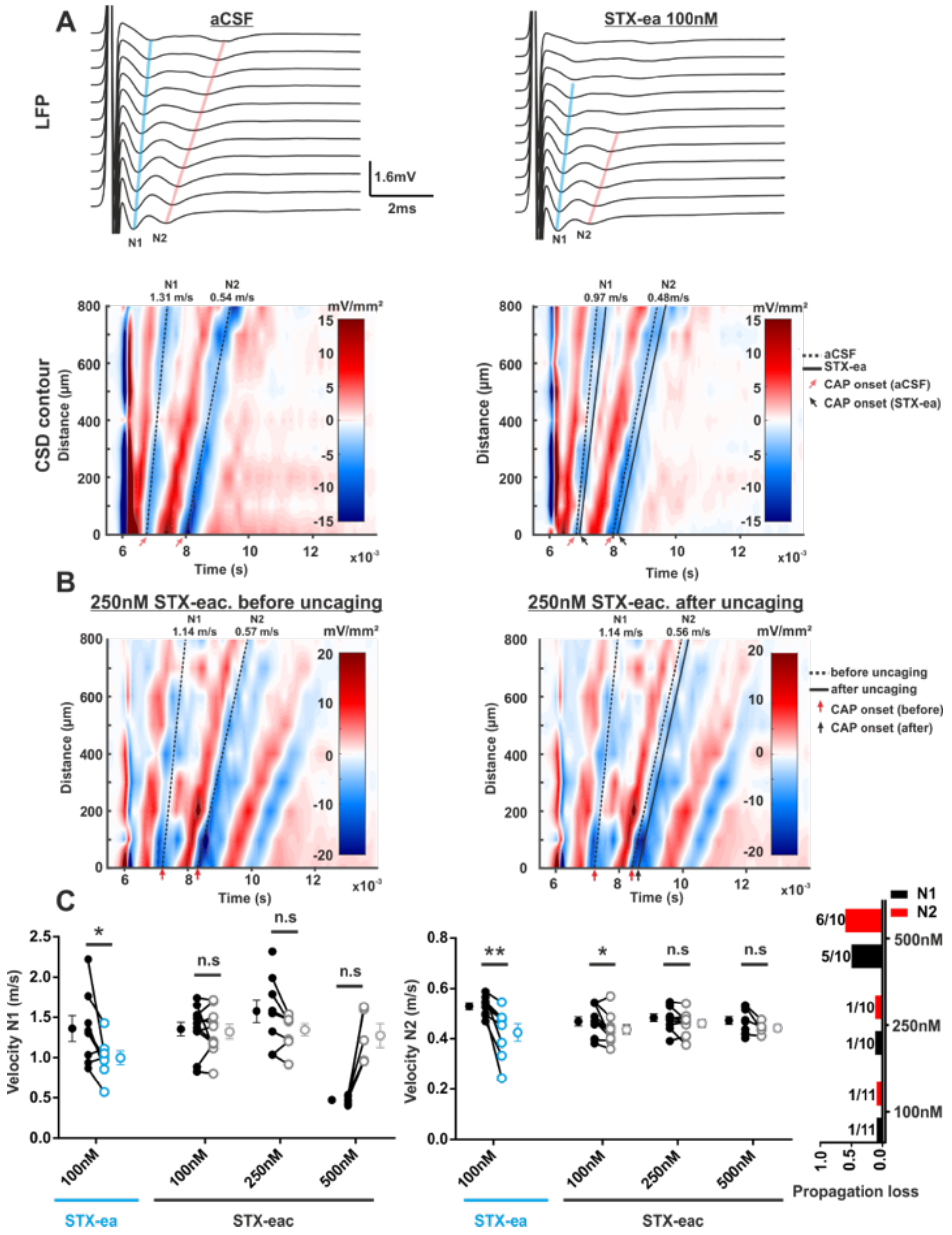


(**A**) Example of LFP signals and CSD contour plot for 100nM **1** applied in bath. The left panel represents the aCSF baseline condition. The right panel represents signals after 10 minutes of application of **1** in bath. For both condition, 10 recordings are averaged. Only the first 11 channels closest to the stimulating electrode are represented. Propagation of N1 and N2 signals across channels is represented respectively in blue and red. In the LFP signal, a reduction of propagation of both N1 and N2 (blue and red traces are shorter) is observed in the 100 nM **1** condition. In the CSD contour plot, both reduction in propagation and delayed onset can be seen. Indeed, after **1** (right, solid lines), there is a shift in both onset and propagation speed compared control (dashed lines). (**B**) Representative CSD contour plots for 100 nM **5** before and after uncaging. The left panel represents uncaged baseline condition. The right panel represents signals after exposure to 500 ms 355 nM light. For both condition, 10 recordings are averaged. Only the first 11 channels closest to the stimulating electrode are represented. Reduction in onset, but not propagation speed is observed for N2, while N1 is apparently unaffected. (**C**) Scatter plot of the effect 100 nM bath application of **1** in blue and of uncaging of 100 nM, 250 nM and 500 nM **5** on the N1 (left) and N2 (middle) velocity. Histograms (on the right) represent the incidence of complete block for N1 (black) and N2 (red) for the three different concentrations of **5**, 100 nM, 250 nM and 500 nM. N1 and N2 velocity are both consistently decreased with bath applied **1**, while little effect was seen with uncaged **5** (except N2 at 100 nM). Notice that half of the slices have a loss of both N1 and N2 propagation at 500 nM, obscuring any potential difference in velocity. Wilcoxon, N=8, 7, 5 and 8 respectively for 100 nM **1**, 100 nM, 250 nM and 500 nM **5**. n.s. $p > 0.05$, *$p < 0.05$, **$p < 0.01$, ***.

#### 2. Extended Data Tables

**Extended Data Table 1.** Uncaging of STX-eac **5** in a corpus callosum slice preparation.

| | Amplitude (mV/mm$^2$) | | | | Time after stimulation (ms) | | | | Velocity (m/s) | | | |
|---|---|---|---|---|---|---|---|---|---|---|---|---|
| | N1 | | N2 | | N1 | | N2 | | N1 | | N2 | |
| | Before | After | Before | After | Before | After | Before | After | Before | After | Before | After |
| **100 nM** | 4.54 ± 1.23 | 4.72 ±1.34 | 5.50 ± 0.75 | 4.1 ± 0.56 | 0.89 ± 0.09 | 0.89 ± 0.08 | 2.47 ± 0.09 | 2.72 ± 0.11 | 1.35 ± 0.08 | 1.32 ± 0.09 | 0.47 ± 0.02 | 0.44 ± 0.02 |
| | n = 11 | n = 11 | n = 11 | n = 11 | n = 11 | n = 11 | n = 11 | n = 11 | n = 11 | n = 10 | n = 11 | n = 10 |
| | p = 0.5117 | | p = 0.0273 | | p = 0.9932 | | p = 0.0039 | | p = 0.9219 | | p = 0.0137 | |
| **250 nM** | 4.98 ± 1.43 | 3.83 ± 1.19 | 5.70 ± 0.80 | 3.66 ± 0.71 | 0.71 ± 0.08 | 0.79 ± 0.05 | 2.36 ± 0.09 | 2.56 ± 0.11 | 1.58 ± 0.14 | 1.35 ± 0.08 | 0.48 ± 0.02 | 0.46 ± 0.02 |
| | n = 10 | n = 10 | n = 10 | n = 10 | n = 10 | n = 9 | n = 10 | n = 10 | n = 9 | n = 8 | n = 10 | n = 9 |
| | p = 0.0156 | | p = 0.0020 | | p = 0.1953 | | p = 0.0117 | | p = 0.1094 | | p = 0.2500 | |
| **500 nM** | 5.59 ± 0.88 | 3.53 ± 0.69 | 6.12 ± 0.99 | 2.56 ± 0.74 | 0.68 ± 0.05 | 0.96 ± 0.06 | 2.40 ± 0.11 | 2.70 ± 0.13 | 1.68 ± 0.19 | 1.27 ± 0.15 | 0.47 ± 0.02 | 0.44 ± 0.01 |
| | n = 10 | n = 10 | n = 10 | n = 10 | n = 10 | n = 9 | n = 7 | n = 7 | n = 9 | n = 5 | n = 10 | n = 4 |
| | p = 0.0191 | | p = 0.0059 | | p = 0.0039 | | p = 0.0625 | | p = 0.1875 | | p = 0.1250 | |

**Before**, before uncaging; **After**, after uncaging.

**Extended Data Table 2.** Bath application of STX-ea **1** in a corpus callosum slice preparation.

| | Amplitude (mV/mm$^2$) | | | | Time after stimulation (ms) | | | | Velocity (m/s) | | | |
|---|---|---|---|---|---|---|---|---|---|---|---|---|
| | N1 | | N2 | | N1 | | N2 | | N1 | | N2 | |
| | aCSF | Bath | aCSF | Bath | aCSF | Bath | aCSF | Bath | aCSF | Bath | aCSF | Bath |
| **100 nM** | 7.44 ± 0.85 | 5.53 ± 1.17 | 12.2 ± 1.97 | 9.07 ± 1.12 | 0.63 ± 0.07 | 0.73 ± 0.08 | 1.91 ± 0.11 | 2.05 ± 0.16 | 1.36 ± 0.16 | 1.00 ± 0.08 | 0.53 ± 0.01 | 0.43 ± 0.03 |
| | n = 8 | n = 8 | n = 8 | n = 8 | n = 8 | n = 8 | n = 8 | n = 8 | n = 8 | n = 8 | n = 8 | n = 8 |
| | p = 0.0156 | | p = 0.0234 | | p = 0.0156 | | p = 0.0078 | | p = 0.0391 | | p = 0.0078 | |

#### 3. Chemical Procedures:

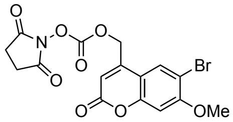


**(6-Bromo-7-methoxycoumarin-4-yl)methoxycarbonyl-N-oxosuccinimide (9).** To a solution of **6** (9.5 mg, 33 µmol) in 666 µL of $CH_3CN$ was added N,N'-disuccinimidyl carbonate (17 mg, 66 µmol, 2.0 equiv) and $Et_3N$ (4.7 µL, 33 µmol). The reaction mixture was stirred for 4 h then diluted with 5 mL of EtOAc and transferred to a separatory funnel containing 5 mL of saturated aqueous $NH_4Cl$. The organic layer was collected and washed successively with 1 x 5 mL of saturated aqueous NaCl and 2 x 5 mL of ½ saturated aqueous NaCl. The organic fraction was dried over $Na_2SO_4$, filtered, and concentrated under reduced pressure to a white powder. Purification of this material by chromatography on silica gel (gradient elution: 19:1→17:3 $CHCl_3$/acetone) afforded **9** (2 mg, 14%) as a white powder.

TLC (9:1 $CHCl_3$/acetone): $R_f = 0.57$

$^1$H NMR (500 MHz, $CDCl_3$) δ 7.64 (s, 1H), 6.89 (s, 1H), 6.47 (s, 1H), 5.43 (s, 2H), 3.98 (s, 3H), 2.88 (s, 4H) ppm

$^{13}$C NMR (125 MHz, $CDCl_3$) δ 168.3, 159.8, 159.2, 154.8, 151.4, 145.6, 127.4, 112.1, 111.0, 108.3, 100.9, 67.0, 57.0, 25.6 ppm

IR (thin film) ν 3093, 2923, 1817, 1791, 1732, 1608, 1413, 1274, 1204, 1080 $cm^{-1}$

HRMS ($ESI^+$): $[MH]^+$ calcd for $C_{16}H_{12}BrNO_8$, 425.9819; found, 425.9808

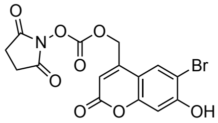


**(6-Bromo-7-hydroxycoumarin-4-yl)methoxycarbonyl-N-oxosuccinimide (10).** This compound was prepared in an analogous manner to **9** starting from **7** (27 mg, 100 µmol) using 2.0 equiv of triethylamine (28 µL, 200 µmol). Purification by chromatography on silica gel (gradient elution: 5:0→4:1 $CHCl_3$/acetone) afforded **5** (25 mg, 60%) as a white powder.

TLC (9:1 $CHCl_3$/acetone): $R_f = 0.32$

$^1$H NMR (500 MHz, $CD_3OD$) δ 7.91 (s, 1H), 6.91 (s, 1H), 6.24 (s, 1H), 5.68 (s, 2H), 2.81 (s, 4H) ppm

$^{13}$C NMR (125 MHz, $CD_3OD$) δ 179.2, 168.8, 167.2, 163.4, 160.4, 157.0, 138.2, 119.5, 119.4, 115.8, 112.7, 76.9, 34.9 ppm

IR (thin film) ν 3088, 1818, 1791, 1738, 1606, 1388, 1275, 1216, 1079 $cm^{-1}$

HRMS ($ESI^+$): $[MH]^+$ calcd for $C_{15}H_{10}BrNO_8$, 411.9663; found, 411.9652

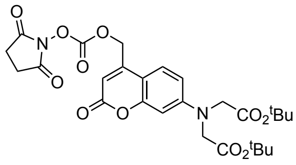

**(7-[Bis(tert-butoxycarbonylmethyl)-amino]coumarin-4-yl)methoxycarbonyl-N-oxosuccinimide (11).** This compound was prepared in an analogous manner to **9** starting from **8** (50 mg, 120 μmol). Purification by chromatography on silica gel (gradient elution: 7:3→1:1 hexanes/EtOAc) afforded **7** (54 mg, 81%) as a yellow oil.

TLC (1:1 hexanes/EtOAc): $R_f = 0.21$

$^1$H NMR (500 MHz, $CDCl_3$) δ 7.30 (dd, *J* = 8.9, 2.2 Hz, 1H), 6.53 (dd, *J* = 8.9, 2.6 Hz, 1H), 6.47 (d, *J* = 2.6 Hz, 1H), 6.29 (s, 1H), 5.40 (s, 2H), 4.05 (s, 4H), 2.86 (s, 4H), 1.47 (s, 18H) ppm

$^{13}$C NMR (125 MHz, $CDCl_3$) δ 168.9, 168.4, 161.0, 155.8, 151.6, 151.4, 146.5, 124.5, 109.4, 109.3, 107.6, 99.4, 82.7, 67.4, 54.4, 28.2, 25.5 ppm

IR (thin film) ν 3467, 2979, 2932, 1791, 1741, 1613, 1394, 1226, 1153 $cm^{-1}$

HRMS ($ESI^+$): $[MH]^+$ calcd for $C_{27}H_{32}N_2O_{11}$, 561.2079; found, 561.2064

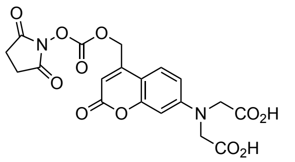


**(7-[Bis(carboxymethyl)-amino]coumarin-4-yl)methoxycarbonyl-N-oxosuccinimide (12).** To a solution of **11** (23 mg, 40.9 μmol) in 0.9 mL of $CH_2Cl_2$ was added 2.8 mL of $CF_3CO_2H$ and 40 μL of $H_2O$. The reaction mixture was stirred for 30 min then concentrated under reduced pressure to afford **12** (25 mg, quantitative) as a yellow foam. This material was used in the subsequent without purification.

TLC (1:1 hexanes/EtOAc): $R_f = 0.00$

$^1$H NMR (500 MHz, $d_6$-DMSO) δ 7.50 (d, *J* = 9.0 Hz, 1H), 6.63 (dd, *J* = 9.0, 2.6 Hz, 1H), 6.51 (d, *J* = 2.6 Hz, 1H), 6.10 (s, 1H), 5.66 (s, 2H), 4.24 (s, 4H), 2.83 (s, 4H) ppm

$^{13}$C NMR (125 MHz, $d_6$-DMSO) δ 171.3, 169.8, 160.13, 155.2, 151.7, 151.0, 148.3, 125.5, 109.3, 107.2, 106.4, 98.1, 67.6, 52.7, 25.4 ppm

IR (thin film) ν 3466, 2941, 1790, 1739, 1612, 1420, 1222 $cm^{-1}$

HRMS ($ESI^+$): $[MH]^+$ calcd for $C_{19}H_{16}N_2O_{11}$, 449.0827; found, 449.0816

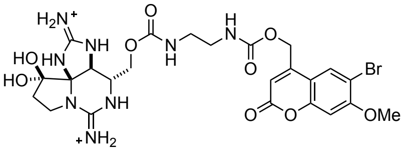


**(6-Bromo-7-methoxycoumarin-4-yl)methyl N21-ethylcarbamate saxitoxin (2).** To an ice-cold solution of STX-ea **1** (1.64 μmol) in 164 μL of pH 8.5 aqueous phosphate buffer (0.1 M $Na_2HPO_4$/$Na_3PO_4$) was added a solution of **9** (1.4 mg, 3.29 μmol, 2.0 equiv) in 164 μL of $CH_3CN$. The reaction flask was stoppered and placed in a sonication bath for 30 seconds. The flask was then wrapped in foil and the contents stirred for 5 h. Following this time, the reaction was quenched by the addition of 16.4 μL of 1.0 M aqueous $CF_3CO_2H$. The reaction mixture diluted with 1.65 mL of a 9:1 10 mM aqueous $CF_3CO_2H$/$CH_3CN$ solution and filtered through a Fisher 0.22 μm PTFE filter. The product was purified by reversed-phase HPLC (Silicycle SiliaChrom AQ C18, 5 μm, 10 x 250 mm column, eluting with a gradient flow of 0→40% $CH_3CN$ in 10 mM aqueous $CF_3CO_2H$ over 80 min, 214 nm UV detection). At a flow rate of 4 mL/min, **2** had a retention time of 47–50 min and was isolated as a white powder following lyophilization (0.48 μmol, 29%, $^1$H NMR quantitation).

$^1$H NMR (600 MHz, $D_2O$) δ 7.98 (s, 1H), 7.17 (s, 1H), 6.39 (s, 1H), 5.35 (s, 2H), 4.71 (s, 1H), 4.28 (dd, *J* = 11.8, 10.0 Hz, 1H), 4.02 (s, 3H), 4.01–3.99 (m, 1H), 3.84–3.76 (m, 2H), 3.57 (dd, *J* = 9.4, 9.4, 1H), 3.40–3.24 (m, 4H), 2.43 (dd, *J* = 14.2, 7.8 Hz, 1H), 2.35 (ddd, 14.1, 9.9, 9.9 Hz, 1H) ppm

HRMS (ESI$^+$): [MH]$^+$ calcd for $C_{24}H_{29}BrN_8O_9$, 653.1314; found, 653.1305

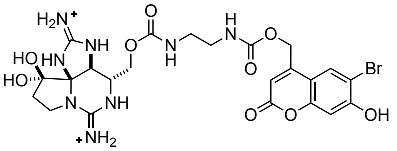

**(6-Bromo-7-hydroxycoumarin-4-yl)methyl N21-ethylcarbamate saxitoxin (3).** This compound was prepared in an analogous manner to **2** starting from STX-ea **1** (2.24 μmol) and **10** (1.8 mg, 4.47 μmol, 2.0 equiv). At a flow rate of 4 mL/min (gradient flow of 0→40% $CH_3CN$ in 10 mM aqueous $CF_3CO_2H$ over 80 min, 214 nm UV detection), **3** had a retention time of 41–42 min and was isolated as a white powder following lyophilization (1.10 μmol, 49%, $^1$H NMR quantitation).

$^1$H NMR (600 MHz, $D_2O$) δ 7.95 (s, 1H), 7.03 (s, 1H), 6.36 (s, 1H), 5.33 (s, 2H), 4.70 (s, 1H), 4.28 (dd, *J* = 12.5, 9.1 Hz, 1H), 4.02–3.96 (m, 1H), 4.83–3.72 (m, 2H), 3.56 (t, *J* = 9.9, 1H), 3.41–3.21 (m, 4H), 2.46–2.39 (m, 1H), 2.38–2.30 (m, 1H) ppm

HRMS (ESI$^+$): [MH]$^+$ calcd for $C_{23}H_{27}BrN_8O_9$, 639.1157; found, 639.1161

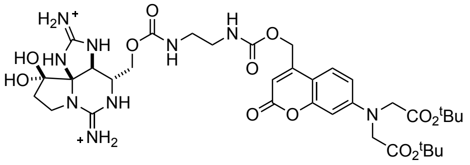

**(7-[Bis(tert-butoxycarbonylmethyl)-amino]coumarin-4-yl)methyl N21-ethylcarbamate saxitoxin (4).** This compound was prepared in an analogous manner to **2** starting from STX-ea **1** (1.73 μmol) and **11** (2 mg, 3.566 μmol, 2 eq). At a flow rate of 4 mL/min (gradient flow of 10→50% $CH_3CN$ in 10 mM aqueous $CF_3CO_2H$ over 80 min, 214 nm UV detection), **4** had a retention time of 54–56 min and was isolated as a white powder following lyophilization (0.29 μmol, 17%, $^1$H NMR quantitation).

$^1$H NMR (600 MHz, $D_2O$) δ 7.60 (d, *J* = 9.0 Hz, 1H), 6.74 (dd, *J* = 9.1, 2.6 Hz, 1H), 6.66 (d, *J* = 2.6 Hz, 1H), 6.23 (s, 1H), 5.40–5.31 (m, 2H), 4.68 (s, 1H), 4.27 (s, 4H), 4.27–4.23 (m, 1H), 3.96 (dd, *J* = 11.8, 5.4 Hz, 1H), 3.77 (t, *J* = 10.0 Hz, 1H), 3.74–3.68 (m, 1H), 3.53 (dd *J* = 9.4, 9.4, 1H), 3.41–3.31 (m, 1H), 3.31–3.18 (m, 3H), 2.44–2.19 (m, 2H), 1.48 (s, 18H) ppm

HRMS (ESI$^+$): [MH]$^+$ calcd for $C_{35}H_{49}N_9O_{12}$, 788.3574; found, 788.3560

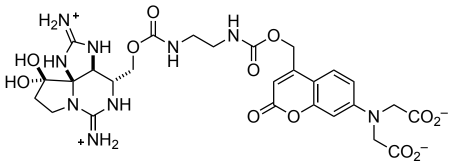

**(7-[Bis(carboxymethyl)-amino]coumarin-4-yl)methyl N21-ethylcarbamate saxitoxin (STX-eac, 5).** This compound was prepared in an analogous manner to **2** starting from STX-ea **1** (3.54 μmol) and **12** (3.3 mg, 7.07 μmol, 2.0 equiv). At a flow rate of 4 mL/min (gradient flow of 0→30% $CH_3CN$ in 10 mM aqueous heptafluorobutyric acid over 60 min, 214 nm UV detection), **5** had a retention time of 44–46 min and was isolated as a white powder following lyophilization (2.17 µmol, 61%, $^{1}$H NMR quantitation). This material was diluted in 9:1 10 mM aqueous $CF_3CO_2H/CH_3CN$ solution and re-lyophilized after quantitation in order to exchange the heptafluorobutyrate counterions for trifluoroacetate counterions.

$^{1}$H NMR (600 MHz, $D_2O$) δ 7.61 (d, *J* = 8.9 Hz, 1H), 6.69 (dd, *J* = 8.9, 2.6 Hz, 1H), 6.58 (d, *J* = 2.6 Hz, 1H), 6.22 (s, 1H), 5.40–5.26 (m, 2H), 4.60 (s, 1H), 4.32 (s, 4H), 4.25 (t, *J* = 10.5 Hz, 1H), 3.91 (dd, *J* = 11.6, 5.3 Hz, 1H), 3.71 (t, *J* = 10.0 Hz, 1H), 3.59–3.53 (m, 1H), 3.51–3.35 (m, 2H), 3.32–3.15 (m, 3H), 2.42–2.24 (m, 2H) ppm

HRMS (ESI$^{+}$): $[MH]^{+}$ calcd for $C_{27}H_{33}N_9O_{12}$, 676.2322; found, 676.2312.

#### 4. $^{1}$H NMR Spectra

**6-Bromo-7-methoxycoumarin-4-yl)methoxycarbonyl-N-oxosuccinimide (9)**

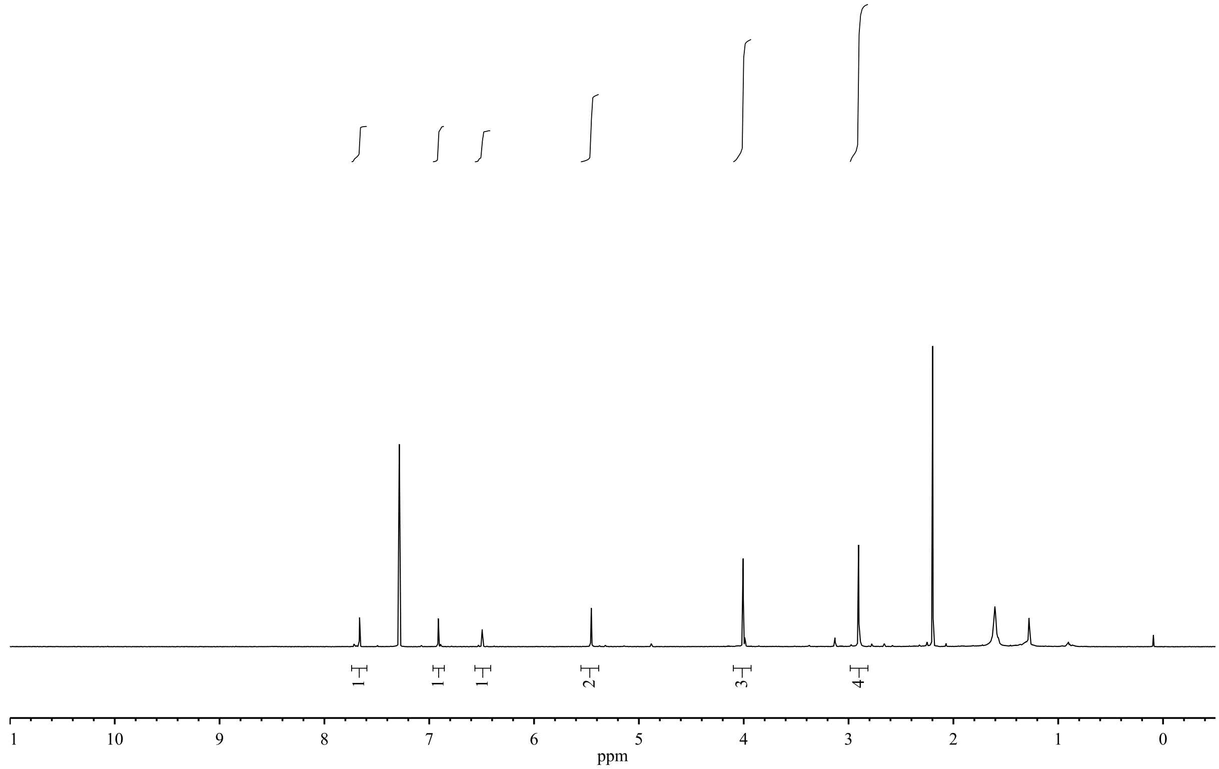


**6-Bromo-7-hydroxycoumarin-4-yl)methoxycarbonyl-N-oxosuccinimide (10)**

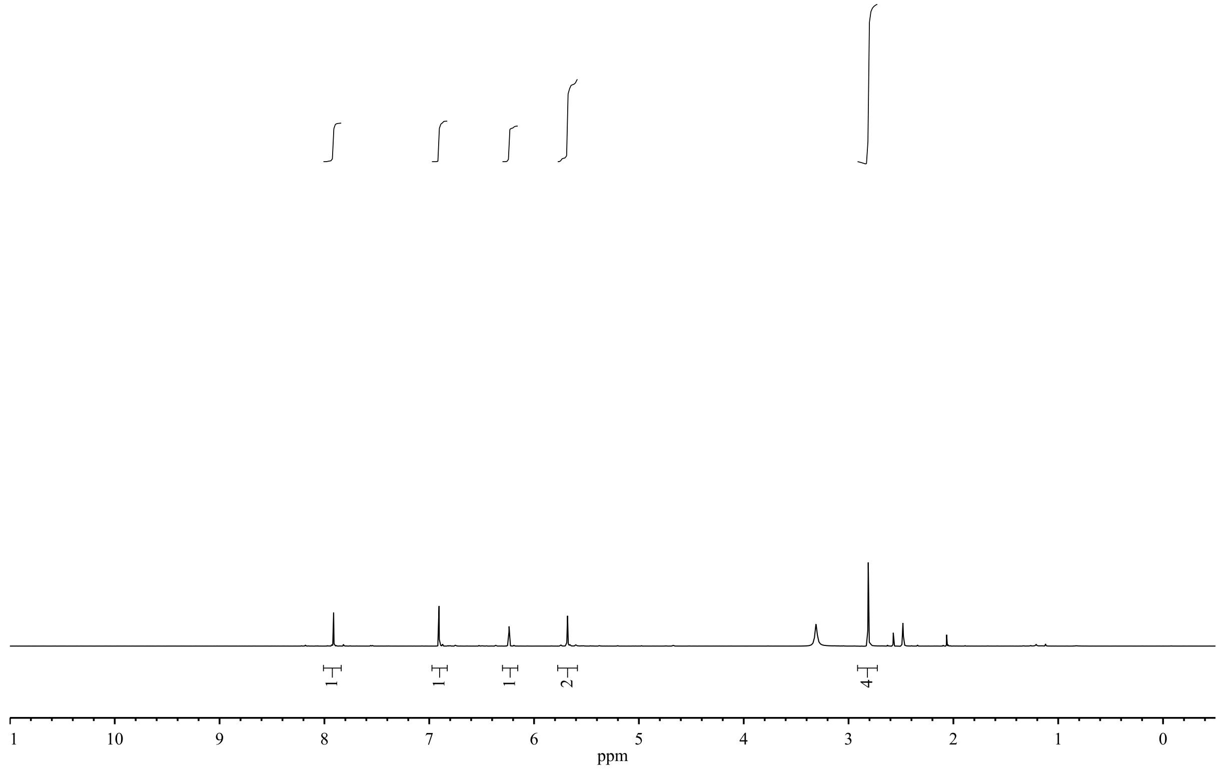

**7-[Bis(tert-butoxycarbonylmethyl)-amino]coumarin-4-yl)methoxycarbonyl-N-oxosuccinimide (11)**

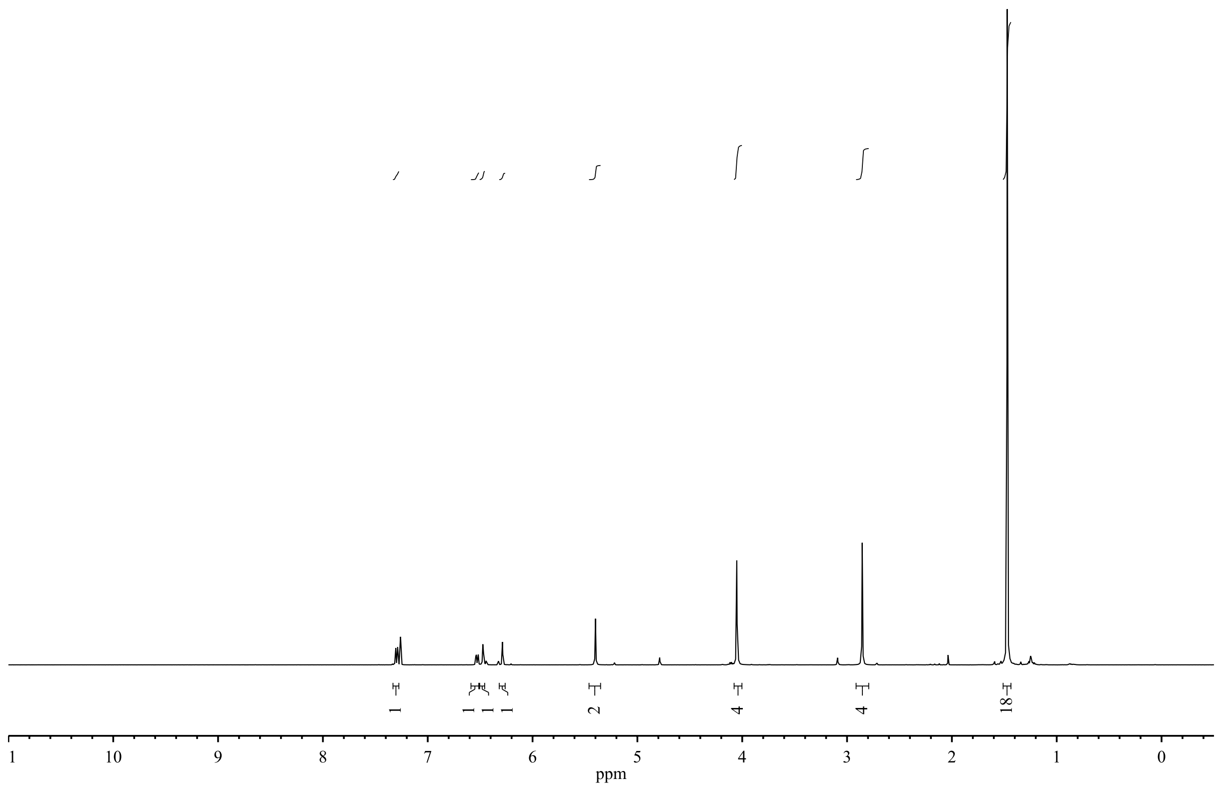


**7-[Bis(carboxymethyl)-amino]coumarin-4-yl)methoxycarbonyl-N-oxosuccinimide (12)**

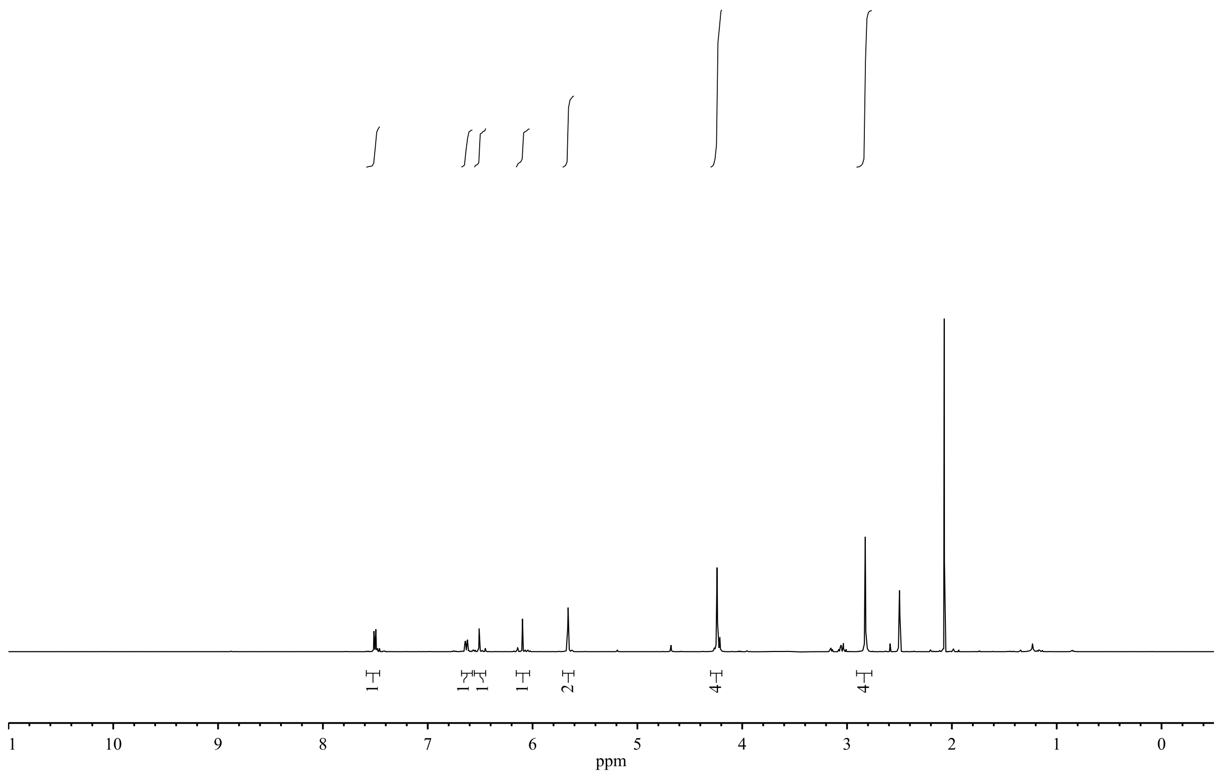

#### 6-Bromo-7-methoxycoumarin-4-ylmethyl N21-ethylcarbamate-saxitoxin (2)

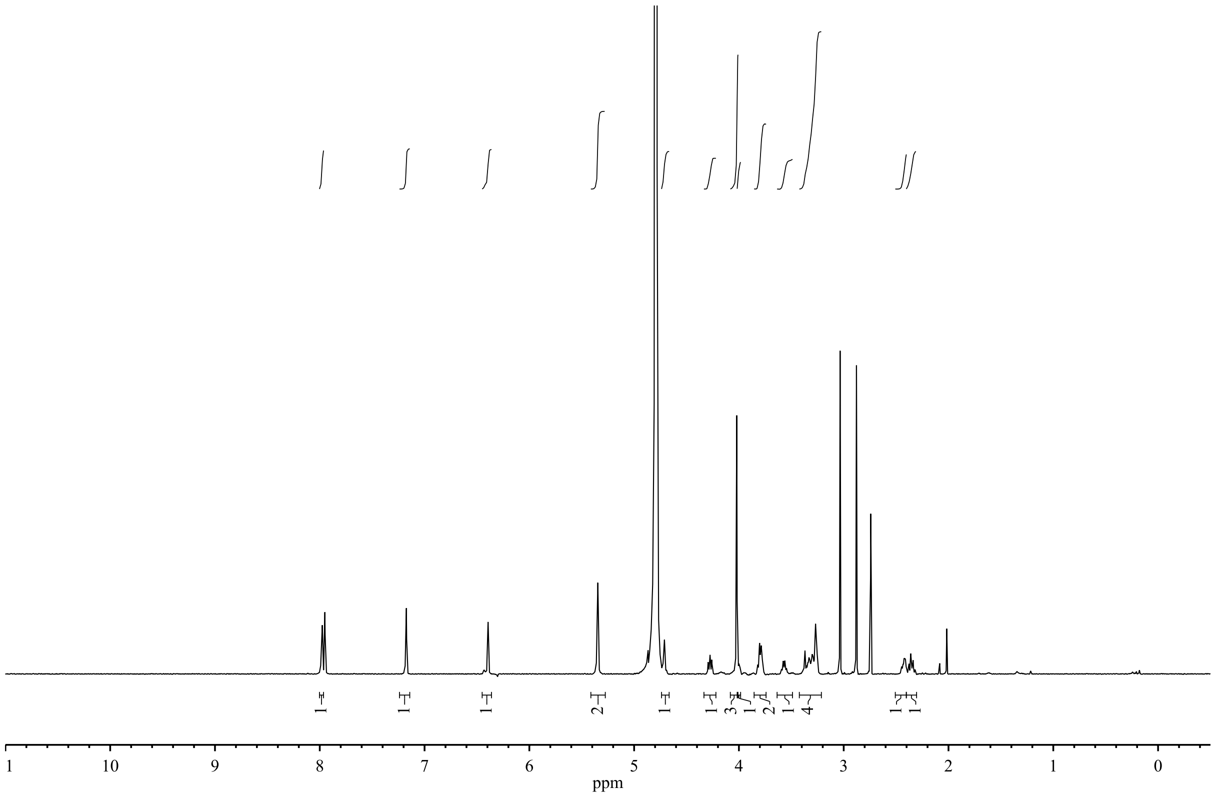


#### 6-Bromo-7-hydroxycoumarin-4-ylmethyl N21-ethylcarbamate-saxitoxin (3)

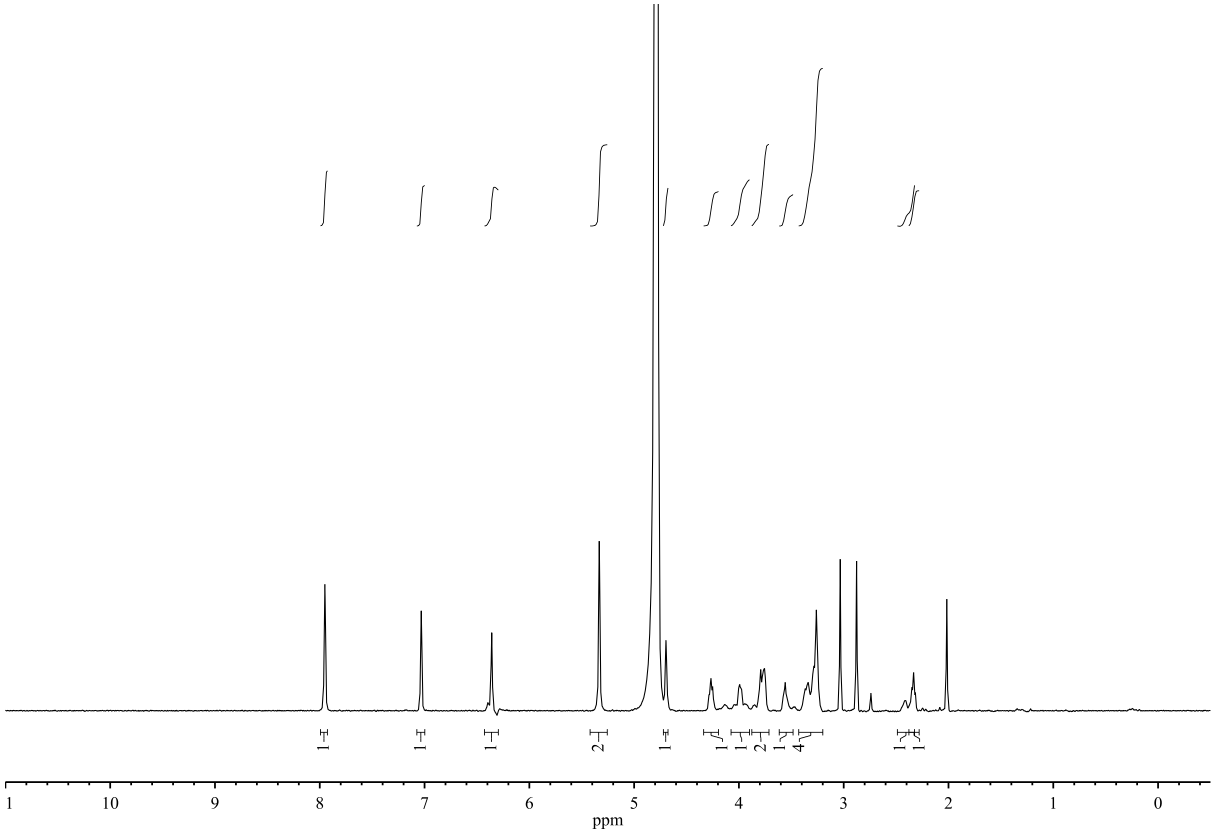

**7-[Bis(tert-butoxycarbonylmethyl)-amino]coumarin-4-yl)methyl N21-ethylcarbamate-saxitoxin (4)**

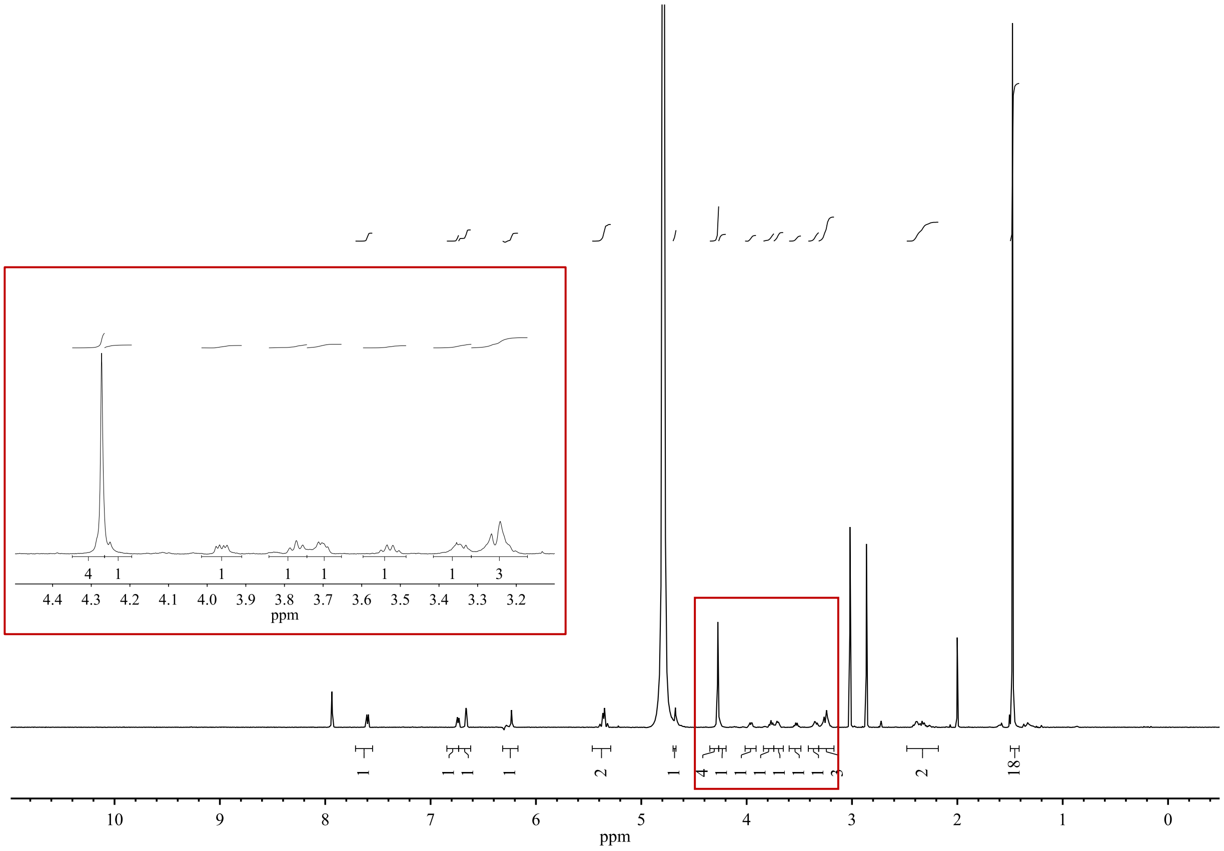

**7-[Bis(carboxymethyl)-amino]coumarin-4-yl)methyl N21-ethylcarbamate-saxitoxin (5)**

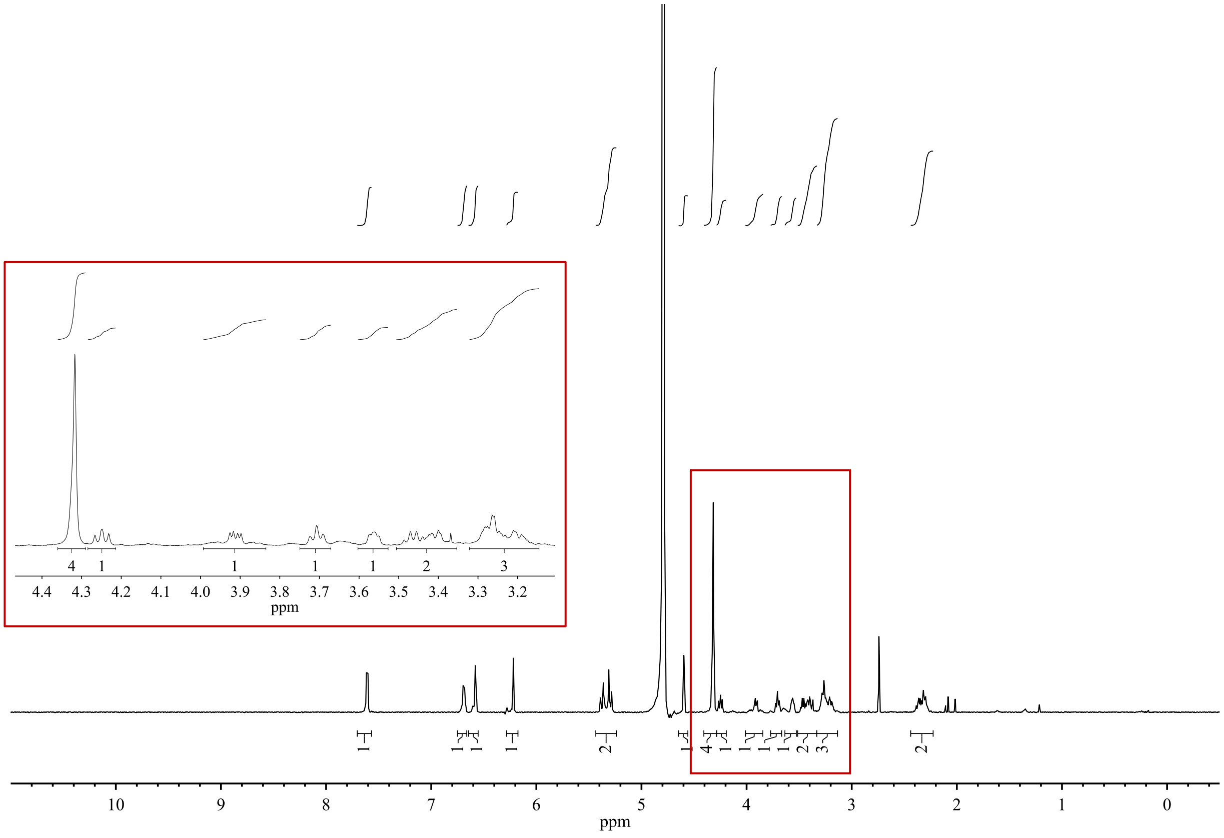

1. Hahn, R. & Strichartz, G. Effects of deuterium oxide on the rate and dissociation constants for saxitoxin and tetrodotoxin action. *J. Gen. Physiol*. **78,** 113–139 (1981).
